# Early differential impact of MeCP2 mutations on functional networks in Rett syndrome patient-derived human cerebral organoids

**DOI:** 10.1101/2024.08.10.607464

**Authors:** Tatsuya Osaki, Chloe Delepine, Yuma Osako, Devorah Kranz, April Levin, Charles Nelson, Michela Fagiolini, Mriganka Sur

## Abstract

Human cerebral organoids derived from induced pluripotent stem cells can recapture early developmental processes and reveal changes involving neurodevelopmental disorders. Mutations in the X-linked methyl-CpG binding protein 2 (MECP2) gene are associated with Rett syndrome, and disease severity varies depending on the location and type of mutation. Here, we focused on neuronal activity in Rett syndrome patient-derived organoids, analyzing two types of MeCP2 mutations – a missense mutation (R306C) and a truncating mutation (V247X) - using calcium imaging with three-photon microscopy. Compared to isogenic controls, we found abnormal neuronal activity in Rett organoids and altered network function based on graph theoretic analyses, with V247X mutations impacting functional responses and connectivity more severely than R306C mutations. These changes paralleled EEG data obtained from patients with comparable mutations. Labeling DLX promoter-driven inhibitory neurons demonstrated differences in activity and functional connectivity of inhibitory and excitatory neurons in the two types of mutation. Transcriptomic analyses revealed HDAC2-associated impairment in R306C organoids and decreased GABA_A_ receptor expression in excitatory neurons in V247X organoids. These findings demonstrate mutation-specific mechanisms of vulnerability in Rett syndrome and suggest targeted strategies for their treatment.

## Introduction

Rett syndrome (RTT) is a progressive neurodevelopmental disorder that is primarily caused by mutations in the X-linked gene encoding methyl-CpG binding protein 2 (MeCP2) and that predominantly affects girls^1,2^. Symptoms typically appear first between 6-18 months of age, and include the loss of purposeful hand skills, decline of speech and social engagement, motor abnormalities, and cognitive impairments^3^. Disease severity can vary significantly among individuals due to the variation of MeCP2 mutations (which include missense and nonsense/truncating mutations) ^4–7^, and while 863 unique mutations have been identfied, eight mutations constitute more than 60% of documented cases^8,9^.

MeCP2 can directly bind DNA selectively through its N-terminal methyl-CpG binding domain (MBD) and recruit other corepressors or coactivators such as Sin3a, HDAC1, HDAC2, or CREB1 through its C-terminal transcriptional repression domain (TRD), resulting in the regulation of a large number of downstream genes ^10,11^. In the past two decades, studies exploring mutations of the MBD (or using MeCP2 null *MeCP2^-/-^* and *Mecp2^-/Y^* mice) have revealed effects of altered MeCP2 affinity to CpG binding regions and their consequences^12–14^. Compared to MeCP2 null and MBD mutations, the impact of the C-terminus is considered to be milder. One such mutation, MeCP2[R306C], in the NCoR-interaction domain (NID) within the TRD, does not affect affinity to the CpG binding domain but abolishes interactions between the NCoR corepressor complex and HDAC3 ^15,16^. Disease severity has also been linked to mutation type in addition to location in the gene^5^. One of the most common truncating mutations in the MBD, MeCP2[R168X], has been associated with severe symptoms^17^. On the other hand, a frequent missense mutation also in the MBD, MeCP2[T158M], has been associated with lesser disease severity ^18,19^. Overall, milder disease severity has been described in Rett patients carrying missense mutations, and the effects of the missense R306C mutation in the NID/TRD have been reported as less functionally deleterious compared to nonsense or truncating mutations in the TRD^20^. Understanding the relationship between mutation types of MeCP2 and disease severity is important for understanding molecular mechanisms underlying the pathology of Rett syndrome as well as developing targeted treatment strategies.

Cerebral organoids derived from human induced pluripotent stem cells (hiPSC) have emerged as powerful experimental systems for their ability to recapitulate early human neurodevelopmental processes^21,22^. This novel *in vitro* approach allows us to expand our current understanding of development which is mostly constructed from studies with human individuals, animal experiments with rodents, and conventional 2D cell cultures *in vitro*. Importantly, patient-derived organoids facilitate analyses of pathological mechanisms underlying not only anatomical alterations^23–25^ but also their neuronal functionality and developmental dynamics^26^. The ability to prepare cerebral organoids from Rett syndrome patients with different mutation types offers the opportunity to directly study the effects of mutations on early developmental events, especially functional networks, and potentially relate disease severity to early occurring misfunction.

In this study, we investigated how two different MeCP2 mutations – the 916C>T / R306C missense mutation and 705delG / V247X truncating mutation, both located near each other in the TRD, with R306C in the NID and V247X just outside it - differentially impact the structure and function of developing neuronal networks. We carried out imaging-based quantification of patient-derived cerebral organoids carrying these two mutations and their isogenic controls using three-photon and two-photon microscopy, employing genetically encoded Ca^2+^ indicators (GCaMP6f and jRGECO1a) along with label-free third-harmonic generation (THG) imaging and additional labeling techniques to identify specific types of neurons. We analyzed the responses and functional connectivity of neurons, including that of inhibitory/excitatory neurons, in MeCP2 mutant and isogenic control organoids and discovered systematic differences in response features between mutation types. Differences in connection patterns were also revealed by network-based analyses based on graph theory. Interestingly, these differences resembled changes observed in EEG data from human patients with comparable mutations, suggesting early developmental origins of brain-wide connectivity. scRNAseq and bulk RNAseq enabled the characterization of different gene expression patterns in the organoids, reinforcing the hypothesis that distinct molecular pathways were affected by different mutations. Specifically, these analyses indicated that inhibitory-to-excitatory connections (GABA-GABRA2/4 and neurexin-neuroligin complex) were affected in V247X organoids whereas HDAC2 gene expression was altered in R306C organoids. These findings suggested mutation-specific strategies for restoring functional responses, and application of specific agents indeed rescued neuronal activity-based connectivity in the two types of organoid.

## Results

### Cerebral organoids derived from Rett syndrome patient-derived iPS cells show mutation-specific structural changes

Cerebral organoids were constructed from Rett syndrome patient-derived iPS cells carrying MeCP2[R306C] and MeCP2[V247X] mutations (**Fig. 1A**). Their isogenic pairs (iPS cells which have the same genetic information except for the MeCP2 mutation) were used as controls (**Fig. S1A, B, C, D**). Isogenic control for the MeCP2[R306C] mutation was created by CRISPR/Cas9 mutagenesis genome editing, and isogenic control for MeCP2[V247X] was cloned and selected from patient iPS cells in which the X chromosome with mutant MeCP2 allele was inactivated (**Fig. 1B, Fig. S1**). Wild-type MeCP2 and mutant MeCP2 expression in these four iPS lines were confirmed by site-specific PCR, Western blotting, and immunostaining (**Fig. S1E, F, and G**) (for detailed differentiation protocols, see Methods and Supplemental Information; **Fig. S2A, B**, and **C**). Before starting differentiation into cerebral organoids, their pluripotency was confirmed by immunostaining with Oct 3/4, Nanog, TRA-1-60, and SSEA4 (**Fig. S1H**). During organoid differentiation (0-3 months), representative iPS markers (Nanog and Oct3/4) decreased, and neuronal progenitor marker (SOX2) and neuron marker (DCX and TUJ1) expression increased (**Fig. S2D, E**). scRNAseq results from 3-month organoids (with both R306C and V247X mutations and isogenic controls) revealed multiple cell types (**Fig. 1C, Fig. S3A**), including neural progenitor cells (expressing *SOX2*), radial glia cells (RGCs, expressing *VIM*), neuronal cells (*DCX*), cycling RGCs (*ASPM*), outer RGCs (oRGCs; *HOPX*) and intermediate progenitor cells (IPCs; *EOMES*). We further identified excitatory neurons (expressing *GRIA2 and SATB2*) and inhibitory progenitors and neurons (expressing *ASCL1, GAD1*, and *GAD2*) as well as astrocytes (expressing *S100B* and *APOE,* **Fig. 1E, Fig. S3C and D**). UMAP plots confirmed multiple cell types in both R306C and V247X organoids (**Fig. 1E**) and highly reduced expression of MeCP2 in V247X organoids, as expected (**Fig. 1D**). MeCP2 was expressed in all cell types (**Fig. S3B**). Cryosectioning showed that the diameter of V247X organoids was significantly larger than isogenic controls throughout the differentiation process, whereas no significant differences between R306C organoids and isogenic controls were observed (**Fig. 1F, Fig. S2B, C**). The number of rosette structures in V247X organoids was higher than in controls, whereas no differences were observed in R306C organoids (**Fig. 1G**). In addition, immuno-histological cryosectioning also revealed structural differences in the neuron layers (**Fig. S2D**): V247X organoids exhibited a thicker postmitotic neuron layer and intermediate layer than isogenic control, whereas R306C organoids did not show significant structural differences (**Fig. 1H)**. After placing cultured organoids onto Matrigel-coated plates, the axonal thickness, migration distance, and axon length were quantified (**Fig. S2F, G, H and I**). V247X organoids showed reduced axon thickness and length, while the R306C organoids showed reduced axon thickness compared to control. These findings demonstrate that a wide range of cell types is present in organoids with both kinds of mutation, and several structural features of organoids show mutation-specific differences.

**Figure 1.**
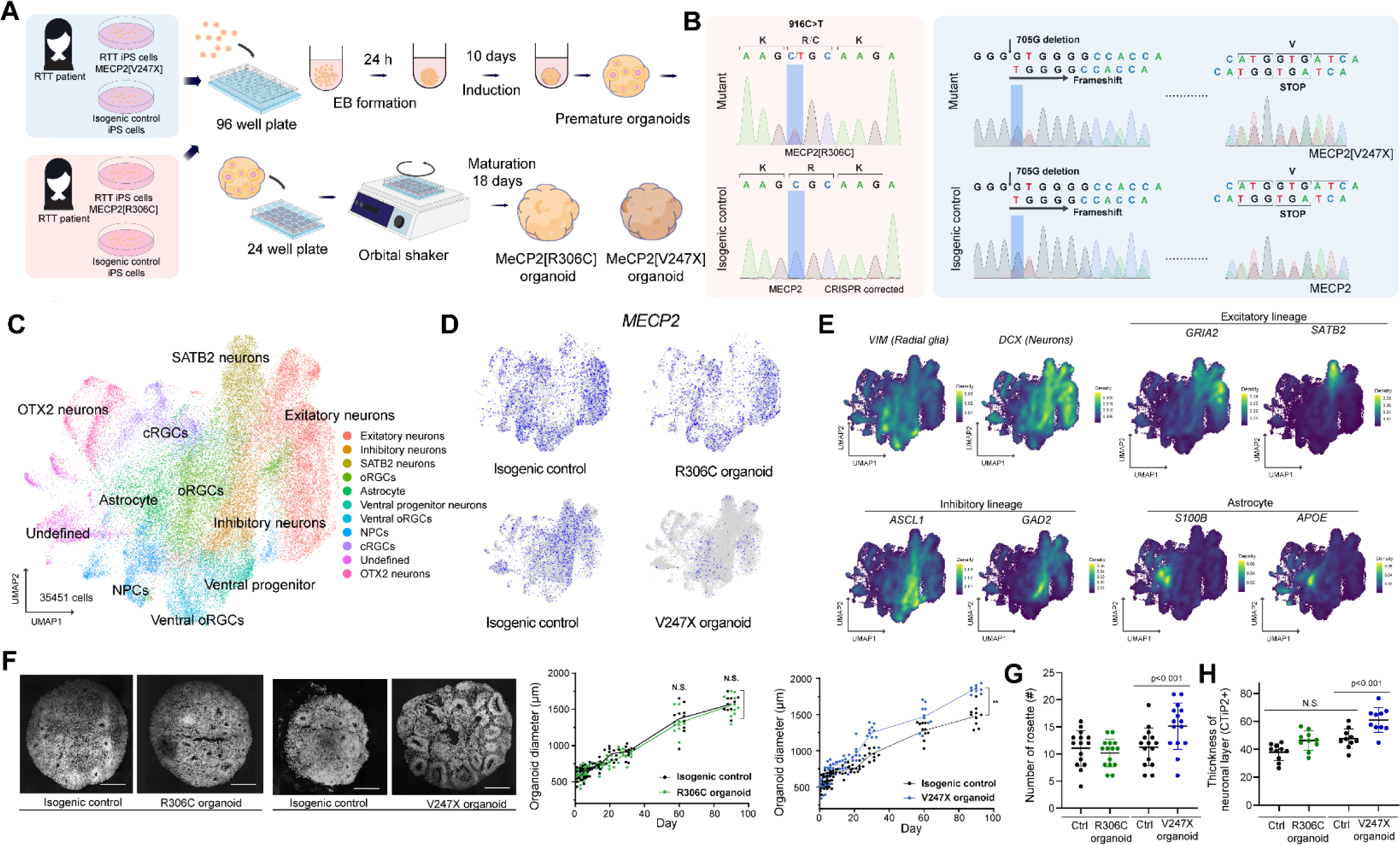
Engineering Rett syndrome patient-derived brain organoids and characterization. (**A**) Schematic illustration of engineering and differentiating brain organoids from human Rett syndrome patient-derived induced pluripotent stem cells (iPS cells). Two different MeCP2 mutations, ([R306C] and [V247X]) and their isogenic controls, were studied. (**B**) Left, MeCP2[R306C]: Sanger sequencing confirmed the heterozygous mutation of 916C>T in MeCP2 and homozygous expression of MeCP2 in both alleles of the X chromosome in isogenic control by CRISPR/Cas9 repairing. Right, MeCP2[V247X]: both mutant and isogenic control have 705G deletion on one of the alleles in the X chromosome, leading to a frameshift mutation; however, the mutant allele in isogenic control iPS cells was inactivated, expressing only wild-type MeCP2. (**C**) Integrated UMAP plot from 4 different types of organoids (MeCP2[R306C] and [V247X] and their isogenic controls). Cells were pooled from 8 individual organoids, analyzing 35351 cells in total. (**D**) Feature plots of MeCP2 expression. V247X organoids expressed no MeCP2 regardless of cell type. (**E**) Feature plots identifying cell types in organoids: radial glia (VIM), inhibitory radial glia (ASCL1), cortical excitatory neurons (DCX, GRIA2, SATB2), inhibitory neurons (GAD2), and astrocytes (S100B and APOE). (**F**) Cryosectioning of brain organoids was used to characterize nuclear staining and quantification of organoid diameter, which increased up to 3 months. n = 12 organoids. (**G, H**) Comparisons of number of rosette structures and thickness of CTiP2 layer by immunostaining. n = 10 organoids. \**p*<0.05, \*\**p*<0.01; one-way ANOVA. Error bars indicate ±SD.

### Altered neuronal activity and synchronicity in Rett syndrome organoids

To examine neuronal activity at signal cell resolution, we utilized CAG-driven GCaMP6f, which was encapsulated with AAV and introduced into the organoids. Subsequently, these organoids were transferred into a custom-made microfluidic device for observation with a three-photon microscope (with imaging at 1300 nm) (**Fig. 2A**). The microfluidic device was designed for fixing the geometrical position of cerebral organoids and a temperature feedback system allowed them to be kept at a steady temperature of 37 L C during imaging (**Fig. S5E**). The field of view, including target neurons in cerebral organoids, was determined based on the depth from the surface and the morphology of neurons identified by third harmonic generation (THG) label-free imaging ^25^ (**Fig. 2B**). Quantification of neuronal activity after deconvolution revealed significantly decreased average activity in R306C organoids compared to isogenic controls (**Fig. 2C, D and E**). In V247X organoids, mean spike frequency decreased slightly but significantly, similar to R306C organoids (**Fig. 2C, D and E**). In addition, pairwise correlations between neurons in organoids were also decreased in both mutations, suggesting that R306C and V247X mutations both had reduced synchronicity (**Fig. 2C, F**). In all conditions, treatment with an antagonist of AMPA receptors (CNQX) and NMDA receptors (AP5) with/without an antagonist of GABA receptors (bicuculline) suppressed neuronal activity(**Fig. 2G**). In contrast, bicuculline addition slightly increased mean spike frequency (**Fig. 2G**). Taken together, these results demonstrate that neuronal activity and neuronal networks in cerebral organoids are supported by glutamatergic and GABAergic synaptic connections, and both MeCP2 mutations influence the mean level and synchronicity of activity.

**Figure 2.**
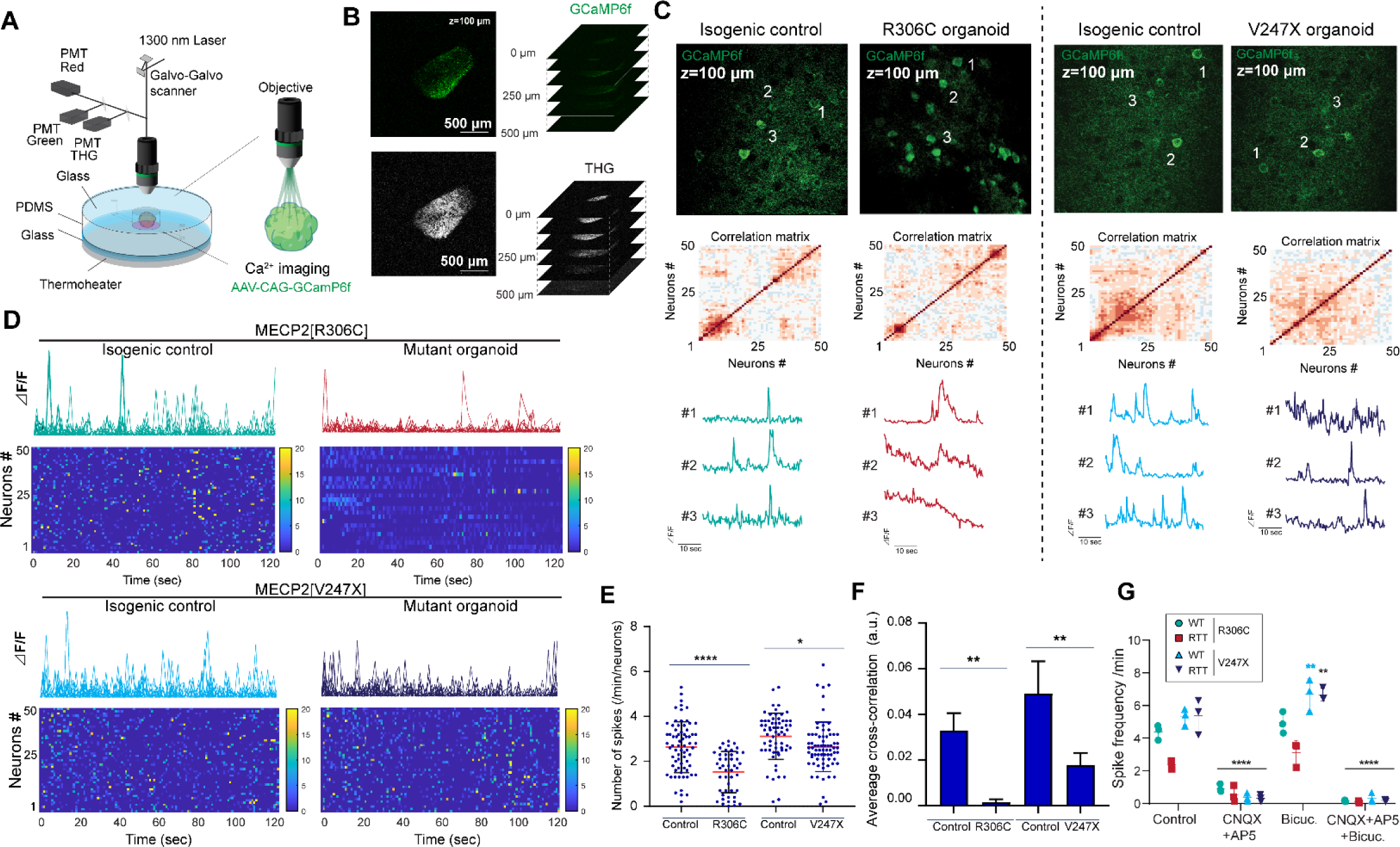
Functional analysis of organoids by Ca^2+^ imaging. **(A)** Brain organoids at 3 months were infected by AAV-CAG-GCaMP6f for Ca^2+^ imaging. Brain organoids were placed in a custom microfluidic device to fix the organoid’s position and prevent movement during the imaging and observation duration. Time series data with Ca^2+^ imaging was taken with a three-photon microscope at 1300 nm by scanning a galvo-galvo mirror. (**B**) Z-slice images of GCaMP6f and label-free Third Harmonic Generation (THG) imaging. THG allows us to determine the structure of organoids. (**C**) Ca^2+^ signals from representative single neurons (n=5 neurons each, at ∼100 um depth) from MeCP2[R306C] and [V247X] organoids and their isogenic control organoids. The correlation matrix shows the synchronized activity among the measured neurons in each field of view (FOV). Both mutant organoids exhibited reduced pair-wise correlation among their neurons. (**D**) Rasters of activity are derived from larger numbers of neurons in each FOV. (**E**) Quantification of spike frequency. Ca^2+^ imaging data was deconvolved to extract spike activity. n = 25. from at least 5 different organoids and 5 different FOV. (**F**) Changes in the average pair-wise correlation between neurons. n = 3. (**G**) Treatment with CNQX and AP5 immediately suppressed the neuronal activity and quantification of spike frequency. Ca^2+^ imaging data in the presence of CNQX and AP5, bicuculine, and all three drugs (CNQX, AP5, bicuculine). Treatment with bicuculine alone slightly but significantly increased spike frequency in MeCP2[V247X] mutant organoids and isogenic controls. n = 3. \**p*<0.05, \*\**p*<0.01; one-way ANOVA. Error bars indicate ±SD.

### Differential impact on functional network architecture in Rett syndrome organoids

To analyze functional network architecture in cerebral organoids, we utilized methods based on graph theory^27^ to examine multi-neuron Ca²L imaging data (see STAR Methods). We hypothesized that the connectivity of neurons might be optimized to mazimize the efficiency of information transfer^28^ even in organoids, and mutant organoids may exhibit inefficient connectivity. To estimate the functional connectivity between neurons and assess their efficiency, we first calculated Pearson’s correlation coefficients between single neuron fluctuations over time (**Fig. 2C, 3A(i)**). Each neuron, represented by its deconvolved ΔF/F Ca^2+^ signal, was treated as a network node. A correlation matrix was constructed by concatenating all pairwise correlation between nodes and thresholded to transform the continuous association matrix into a binary adjacency matrix (**Fig. 3A(i)**) ^27^. Then, graph network models were constructed in two ways: (a) a structural network that retained the spatial information of neurons (**Fig. 3A(ii)**), and (b) a functional network that emphasized topological properties derived from pairwise correlations (such that neurons with higher correlations were situated closer together, leading to clustering), abstracting away from the physical arrangement of neurons to focus on interactions^29, 30^ (**Fig. 3A(iii)**).

**Figure 3.**
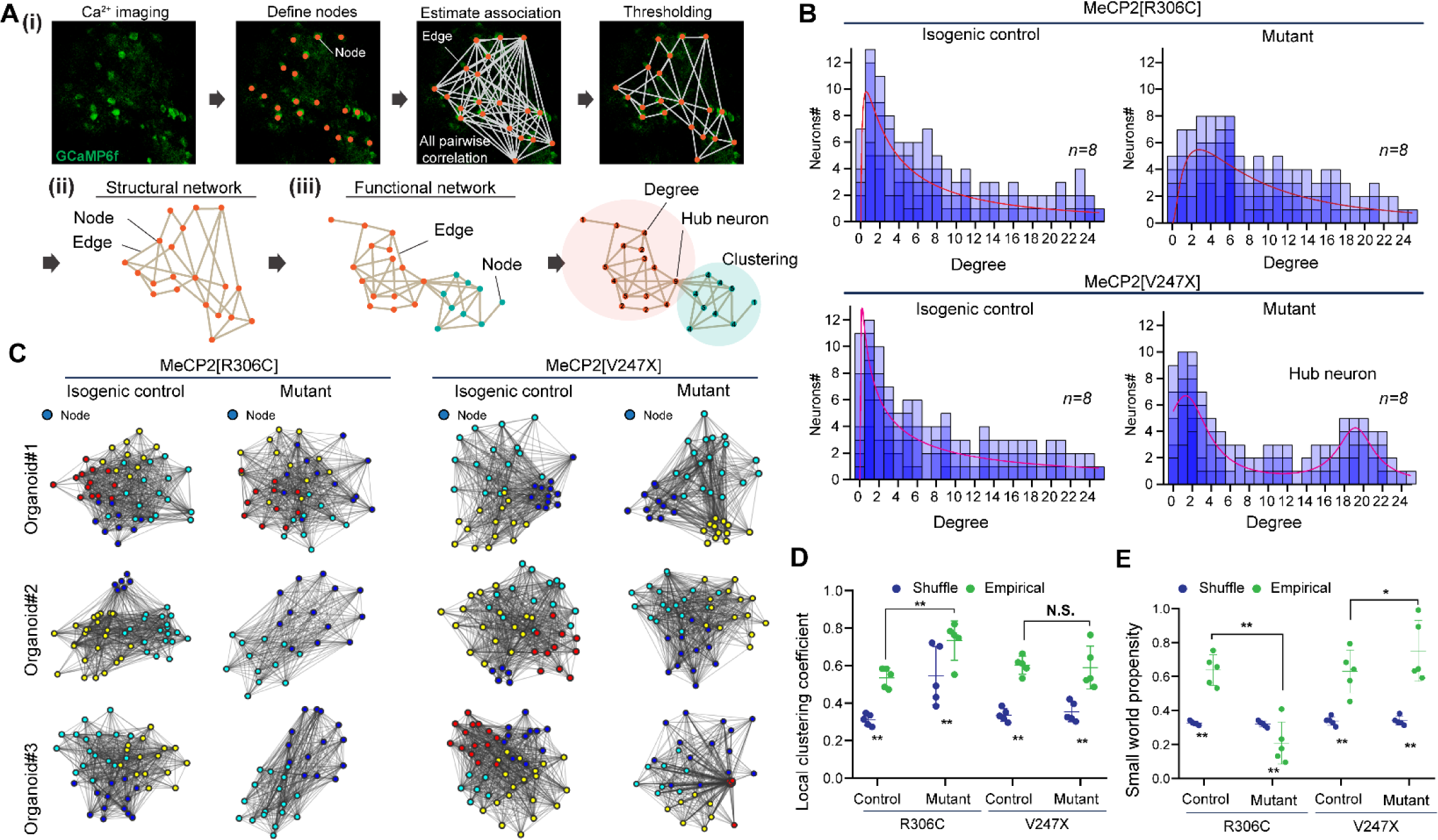
Network analysis of organoids. **(A)** Scheme of functional and structural topological analysis based on graph theory. Individual neurons were identified as nodes, nodes were connected to form edges according to the pairwise correlation between neurons, the connections (edges) were thresholded, and the results were plotted with geometric or neuron location information (as ‘structural network’) or with topological information (as ‘functional network’). The connection degree, hub neurons, local clustering coefficient, and path length were characterized. (**B**) Degree distribution of MeCP2 mutant [R306C] and [V247X] and isogenic control organoids. Degree distributions were fitted with a chi-square curve in R306C and both sets of control organoids. The V247X organoid distribution was fitted with a sum of two Lorentzian distributions. n = 8. (**C**) Plots of functional network maps in topological space; three representative organoids (of n = 8). Colored nodes represent neurons that belong to the same cluster. (**D**) Quantified result of local clustering coefficient. ‘Empirical’ denotes data from actual time series used for correlation calculation. ‘Shuffle’ denotes randomly shuffled data as control. (**E**) Small world propensity calculated using the local clustering coefficient and path length. \**p*<0.05, \*\**p*<0.01; one-way ANOVA. Error bars indicate ±SD.

First, we counted the number of connections for each node (**Fig. 3A, B**) (referred to as its “degree”, with the distribution of the degree across all nodes referred to as the “degree distribution”). The degree distribution for different organoids indicated distinct connectivity profiles, indicating differences in the extent to which neurons were connected to neighboring neurons. For R306C organoids, the amplitude and geometric mean of the distribution (fitted with a chi-square distribution) exhibited a higher value compared to controls, indicating a greater number of connections per neuron. For V247X organoids, the amplitude and geometric mean of the distribution were small and similar to controls. However, a few nodes had a very large degree, indicating the existence of so-called “hub” neurons that made significantly more connections than the others^31,32^. The distribution (fitted with a sum of two Lorentzian distributions) reflected a significant number of highly connected hub neurons. The architecture of the functional network across recorded neurons for R306C and V247X organoids (**Fig 3C**) was used to derive the local clustering coefficient, which was larger in R306C organoids than in isogenic controls (comparing ‘empirical’ or experimentally observed coefficients with ‘shuffled’), indicating more local clustering of coactive neurons (**Fig. 3D**). For V247X organoids, in contrast, no significant difference in local clustering coefficient was observed compared to controls. To obtain a quantitative measure of functional network architecture, we calculated their small-world propensity (SWP), a parameter combining local clustering with path lengths: high SWP relative to shuffled indicates higher clustering and shorter path length, and is usually consistent with more efficient information segregation and integration^27^. Our results showed that both R306C and V247X mutations affected small-world propensity – SWP was significantly decreased in R306C organoids compared to controls but slightly increased in V247X organoids, whereas SWPs in isogenic controls were equivalent (**Fig. 3E**).

Each organoid rosette and its associated ventricular zone may reflect the precursor of a cerebral hemisphere^33^. SWP is a measure of network organization that can be applied across scales^27^. Thus, we next asked whether alterations of network structure in organoids represented changes in cortical organization that could also be discerned in human Rett syndrome patients. EEGs were recorded from the scalp surface of five Rett syndrome patients with two types of truncating mutations in the TRD, MeCP2[R255X] and [R270X], and the missense MeCP2[R306C] mutation in the NID, along with age-matched controls (**Fig. S4**). EEG data were analyzed with graph theoretic approaches based on wavelet coherence with specific frequency windows (**Fig. S4A, B, and C**). Power spectrum density (PSD) analyses from the Rett syndrome patients showed variable changes across frequency bands in the cases with R255X and R270X mutations (**Fig. S4C, D**). In the case with R306C mutation, PSD in beta-band frequency was significantly lower than in the control, consistent with previous findings^34^. Similar to organoids, we analyzed the connectivity between different brain regions based on graph theory (**Fig. S5A, B, and C**). Cases with R255X and R270X mutations showed roughly similar correlations between multiple brain regions compared to controls, whereas brain activity in the Rett patient with R306C mutation appeared to be more strongly correlated across multiple brain regions compared to control (**Fig S5A**). In addition, network structure defined by SWP (**Fig. S5C**) showed that SWP values of patients with truncating mutations were slightly decreased compared to controls (R255X mutation) or slightly increased (R270X mutation). In contrast, SWP values in the patient with R306C mutation were strongly decreased compared to control. This trend is consistent with the data from brain organoids with the R306C mutation (**Fig. 3E**). While based on a limited number of subjects, these preliminary analyses suggest that some large-scale patterns of connectivity changes discerned in Rett syndrome patients may have their roots in early stages of development.

Finally, we examined in organoids whether the disposition of connections (i.e., their spatial orientation or direction) was affected by RTT mutations. Three-photon simultaneous Ca^2+^ imaging and label-free THG imaging enabled the determination of layer structures within organoids and the measurement of neuronal activity across the superficial, intermediate, and SVZ/VZ layers (**Fig. 4A**). This layer-specific information was subsequently incorporated into the analysis of neuronal activity based on Ca^2+^ imaging. The analyses revealed significant suppression of neurons in the intermediate layer in both kinds of mutant organoids compared to isogenic controls, with no significant differences observed in the neurons of the superficial and SVZ/VZ layers (**Fig. 4B**). Additionally, we examined whether inter-neuronal correlations were differentially influenced by the location of neurons within layers or across inter-layer zones. We calculated pairwise correlations and related them to the angle (θ) formed by the line connecting the two neurons and the line connecting one neuron and the center of rosette-like structures (thus, θ values would be high for intra-layer pairs and low for inter-layer pairs, and cos θ would vary from 0 to 1 respectively) (**Fig. 4C**). Our findings indicate that neurons within the same layer tended to exhibit stronger connectivity than those in different layers across all conditions. However, these trends became less pronounced in RTT mutant organoids, particularly in the case of the V247X mutation (**Fig. 4D**). Thus, RTT mutations alter the spatial disposition of neuronal connections during development in addition to their number, strength and small-world propensity.

**Figure 4.**
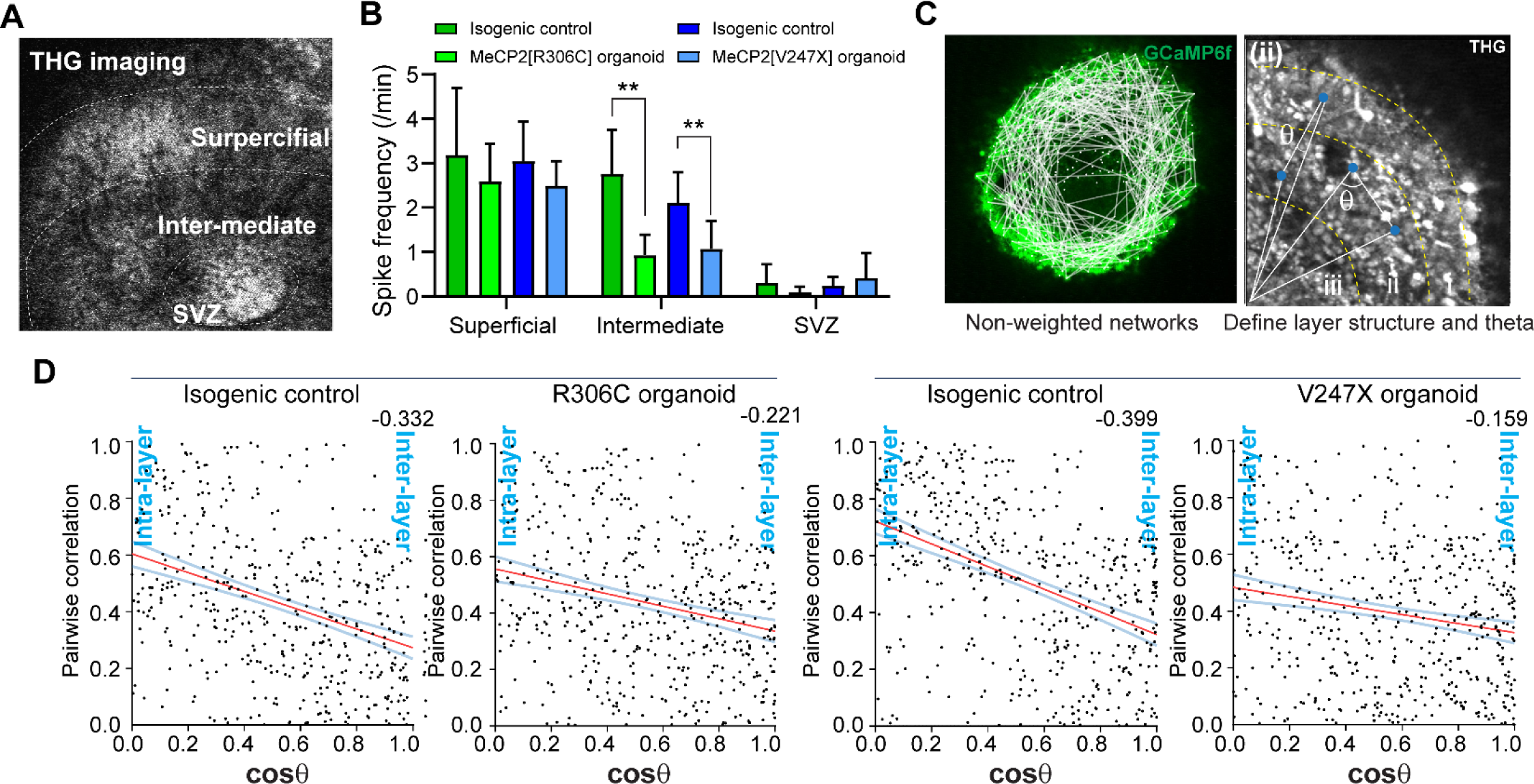
Laminar connectivity analysis of neurons in organoids. **(A)** THG imaging facilitates determination of the layer structure in organoids. (**B**) Spike frequency in each layer was separately calculated based on categorical information about the layer. (**C**) Conceptual images for defining the layer structure by THG imaging and inter/intra layer connection. by calculating the angle (θ) toward the edges from the center of the VZ layer. (**D**) Relationship between angle between neurons and their pairwise correlation. Neurons in the same layer tended to be more connected than in different layers in all conditions; however, these trends become weaker in RTT mutant organoids, especially MeCP2[V247X] mutant organoids. n = 4, pooled from individual four organoids. \**p*<0.05, \*\**p*<0.01; one-way ANOVA. Error bars indicate ±SD.

### Altered gene expression in RTT organoids and the impact of specific genes in R306C and V247X mutations

Changes in functional networks and architecture following Rett mutations may reflect differential gene expression patterns in mutant organoids compared to controls. To identify such changes and their potential impact, we performed both single cell RNAseq (scRNAseq) and bulk RNAseq on organoids isolated from 3-3.5 month cultures (**Fig. 5**). UMAPs from organoids with the R306C mutation and isogenic controls specified multiple subtypes of neurons, as well as radial glia and astrocytes (**Fig. 5A**). We identified excitatory neurons which express FOXG1 and FOXP2, as well as vGLUT and CTiP2. In addition, we confirmed inhibitory lineage cells, which express ASCL1 and GAD2 (**Fig. 5A**). The volcano plot from bulk RNAseq showed 343 upregulated genes and 107 downregulated genes compared to isogenic control organoids (**Fig. 5B**). One of the highest upregulated genes in R306C mutant organoids was *HDAC2 (*Histone Deacetylase 2), one of the co-repressors of MeCP2. GO enrichment analysis revealed that the R306C mutation influenced pathways related to alteration of cellular components such as receptor complex, neuron spine, and dendritic spine, as well as peptidyl-tyrosine modification and biogenesis including proliferation and differentiation (**Fig. S6A**). scRNAseq revealed that the expression level of HDAC2 tended to be altered in relatively mature excitatory and inhibitory neurons and astrocytes, not radial glial cells, even though all cell types expressed HDAC2 (**Fig. 5C**). To infer neuron-neuron communication with the scRNAseq dataset, NeuronChatDB ^35,36^ was employed and predicted altered pre-post synaptic connectivity, especially among excitatory neurons, inhibitory neurons and astrocytes along with decreased number of links between the cells and strength of connections (**Fig. 5E**). Importantly, cell-adhesion complexes which support synaptic connections, especially components of the neurexin–neuroligin signaling pathway (e.g. *NRXN2-NLGN4X, NRXN2-NLGN1, NRXN1-NLGN4X*) and glutamate-glutamate receptor pathway, were deficient in mutant R306C organoids compared to control organoids (**Fig. 5F**).

**Figure 5.**
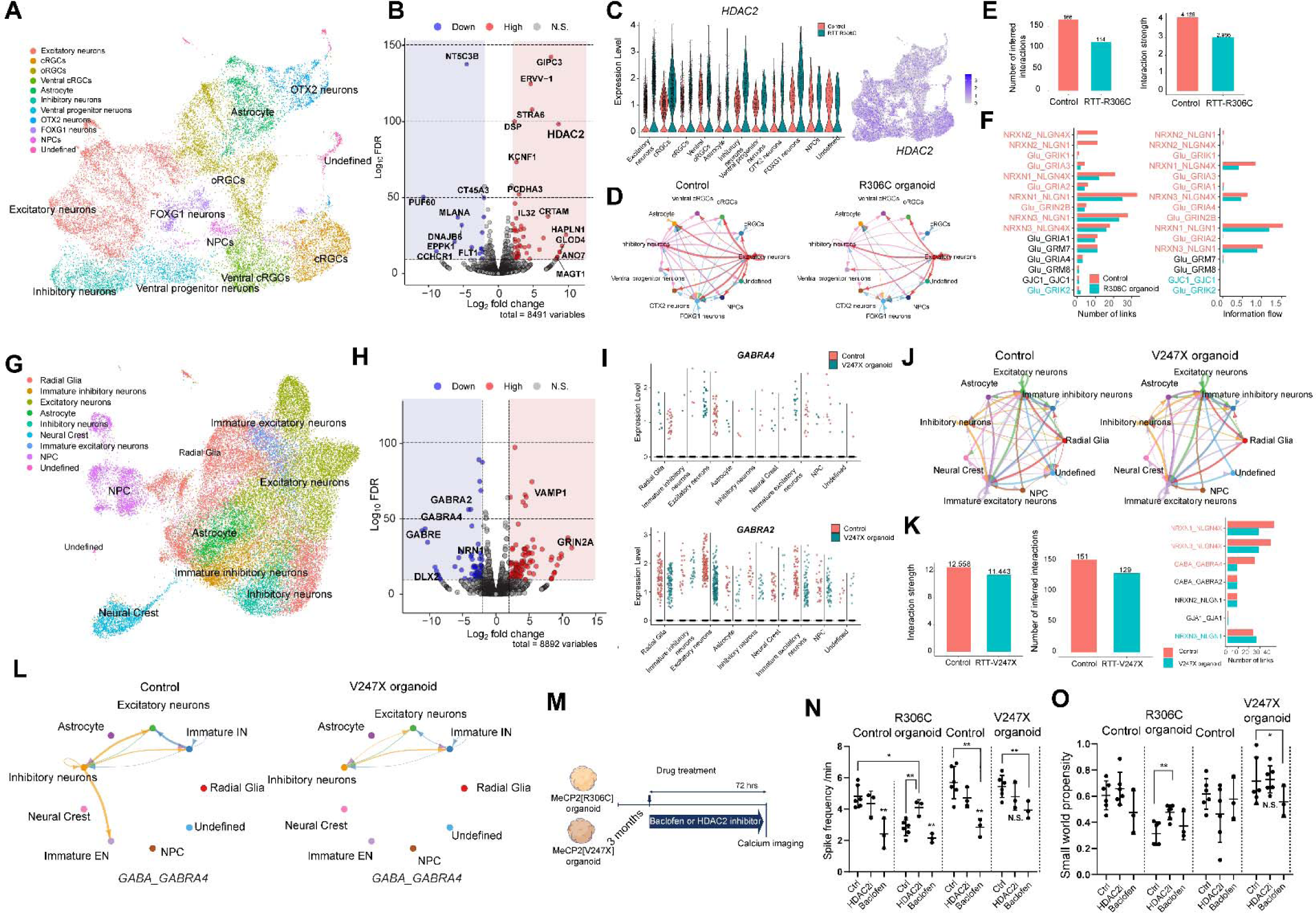
Differential gene expression in mutant organoids defined by bulk RNAseq and scRNAseq. **(A)** Integrated UMAP plot of 3-month cultured RTT-R306C cerebral organoid and isogenic control, identifying the cell types of organoids. (**B**) Differential gene expression in brain organoids carrying MeCP2[R306C] mutation, identifying HDAC2 overexpression in the mutant organoids. RNA was extracted and pooled from 8 different 3-month organoids in each condition for bulk RNAseq. The volcano plot shows 343 upregulated and 107 downregulated genes from bulk RNAseq. (**C**) HDAC2 expression after classifying cell types with scRNAseq. Altered HDAC2 gene expression was observed in all neural cells including excitatory and inhibitory neurons, radial glia, and astrocytes. (**D**) Neuron-neuron connectivity inferred by NeuronChatDB in R306C mutant and control organoids. (**E**) Quantification of differential inferred neuron-neuron interaction. (**F**) Significant downregulated/upregulated signaling pathway among cell types. Downregulated neurexin-neuroglin and Glu-GluR interactions were observed in the R306C mutant organoids. (**G**) Integrated UMAP plot of 3-month cultured RTT-V247X cerebral organoid and isogenic control. (**H**) Differential gene expression in brain organoids carrying MeCP2[V247X] mutation, identifying downregulation of GABAergic receptors in mutant organoids. RNA was extracted and pooled from 6 different organoids in each condition for bulk RNAseq. The volcano plot shows 896 upregulated and 494 downregulated genes. (**I**) GABRA4 and GABRA2 expression in different cell types; downregulation was observed in excitatory but not in inhibitory neurons. (**J**) Inferred neuron-neuron connectivity determined by NeuronChatDB in RTT-V247X mutant and control organoids. (**K**) Quantification of differential inferred neuron-neuron interaction. Downregulated neurexin-neuroglin and GABA-GABRA4 interactions were observed in the RTT-V247X mutant organoids. (**L**) Inferred connectivity analysis showing loss of GABA->GABAR4 directional interaction between excitatory neurons and inhibitory neurons, due to the downregulation of GABARA4 in excitatory neurons. (**M, N, O**) Drug treatment (CAY10683 (HDAC2 inhibitor) and Baclofen) for 72 hrs for cerebral organoids, showing (**N**) spike frequency and **O**) small world propensity in the presence of drugs. n = 3 organoids per condition. \**p*<0.05, \*\**p*<0.01; students’s t-test for two-compasiton and one-way ANOVA for multiple comparison. Error bars indicate ±SD.

In MeCP2[V247X] organoids, scRNAseq revealed fewer diverse and immature neuron and astrocyte subtypes compared to MeCP2[R306C] organoids and isogenic controls (**Fig. 5G**). This lack of diversity and maturity is consistent with immunostaining results (**Fig. 1**), which suggest donor-dependent differences in organoid development. As in R306C organoids, we found excitatory neurons which express FOXG1 and SATB2, and inhibitory lineage cells which express ASCL1 and GAD2. The mutation impacted a large number of genes, with 494 downregulated genes and 896 upregulated genes as shown in the volcano plot (**Fig. 5H**), influencing axon development, regulation of neuron projection development as well as the Wnt signaling pathway (**Fig. S6B**). Highly downregulated genes included genes in the GABA-A receptor family (*GABARA2* and *4*, and *GABARE*), while highly upregulated genes included *GRIN2A* encoding the GluN2A NMDA receptor subunit (**Fig. 5H**). scRNAseq further revealed the cell type-specific down-regulation of *GABRA2* and *GABRA4* (**Fig. 5I**), and upregulation of *GRIN2A* (**Fig. S6C**): these genes were regulated in only excitatory or immature excitatory neurons but not inhibitory neurons. NeuronChatDB^36^ was used to identify the loss of synaptic communication in mutant organoids (**Fig. 5J**), indicating that the number of links and strength of several neurexins–neuroligin and GABA-GABA receptor complex pathways were decreased (**Fig. 5K**). In particular, *NRXN1-NLGN4X*, *NRXN1-NLGN3X*, and *GABA*-*GABRA4* interactions were altered (**Fig. 5K, S6D**). The chord diagram revealed that *GABA* (pre-synaptic in inhibitory neurons)-*GABRA4* (post-synaptic in excitatory neurons) became weaker in V247X mutant organoids over control (**Fig. 5L**).

Analysis of cell-cell communication using the scRNAseq data set with CellChatDB^35^ showed that the total number of interactions in all the cell types was decreased in R306C organoids compared to control, but not in V247X organoids (**Fig. S6E, F**). The strength of communication was slightly decreased in both cases (**Fig. S6G**). We found that astrocytes played an important role in this dysregulation (**Fig. S7**). CellChatDB analysis of differential interactions (**Fig. S7A-C**) showed that in R306C organoids, most intercellular communication was downregulated, and a few (NPC to oRGCs, ventral progenitor cells to ventral oRGCs, and ventral oRGCs to inhibitory neurons) were upregulated. In contrast, astrocyte-to-excitatory neuron and astrocyte-to-immature excitatory neuron communication in V247X organoids were selectively deficient (**Fig. S7B**). In particular in V247X organoids, NRG3 (astrocyte)-ERBB4 (excitatory, astrocyte and radial glia, and NPC), NRXN3 (astrocyte)-NLGN1 (excitatory neurons, immature excitatory neurons), and Glu-SLC1A2/3-GLS to GRIAs intercellular communication were significantly reduced, but not in R306C organoids (**Fig. S7D-G**). Both NRG3-ERBB4 and Glu-SLC1A2/3-GLS to GRIAs signaling pathways are fundamental functions of astrocytes, necessary for forming astrocyte-neuron adhesion and regulating glutamate uptake at glutamatergic synapses^37–39^. Thus, these analyses suggest mutation-specific roles for astrocytes in the pathophysiology of Rett syndrome.

The neuronal gene expression findings demonstrating HDAC2 upregulation in R306C organoids and GABA-A receptor downregulation in V247X organoids led us to examine whether HDAC2 inhibition or GABA activation would rescue neuronal activity or connectivity in these mutant organoid types. Thus a selective inhibitor of HDAC2 (CAY10683) and a GABA agonist (Baclofen) were administrated to 3-month cultured organoids for 72 hours (**Fig. 5M**). The HDAC2 inhibitor increased neuronal activity and restored small-world propensity to control levels in R306C organoids without causing any change in V247X organoids (**Fig. 5N, O**). Baclofen treatment generally reduced the level of neuronal activity in all organoid types (**Fig. 5N**). In V247X organoids, it also reduced small-world propensity, restoring it to control levels, while it caused no change in R306C organoids (**Fig. 5O**). Thus, specific gene expression changes may underlie neuronal activity levels and functional architecture in different RTT mutations, and selective application of molecules that counter these changes may be plausible mechanisms for restoration of function

### Co-DLX labeling with Ca^2+^ imaging to identify functional connectivity in excitatory and inhibitory neurons

The transcriptome analysis above revealed that V247X mutant organoids displayed significant alterations in the development of GABAergic synaptic connectivity. To follow up these findings, GABAergic interneurons expressing DLX were dual-labeled with EGFP and a red Ca^2+^ tracer (jRGECO1a). This dual labeling enabled a comparative assessment of the activity of excitatory and inhibitory neurons individually, as well as their functional connectivity (**Fig. 6A-C, Fig. S8E**). First, flow cytometry and cell sorting were performed to confirm the population of DLX-positive cells, followed by gene expression analysis (**Fig. 6A, Fig. S8A-D**). Flow cytometry and imaging-based counting analyses revealed that in all types of organoids, DLX-positive cells constituted approximately 15% of the total cells, indicating the presence of GABAergic interneurons (**Fig. S8C**). There were no significant differences in the ratio of excitatory to inhibitory neurons among the different types of organoids. The remaining 85% of cells were considered as putative excitatory neurons. Utilizing functional connectivity information from DLX labeling allowed us to define the node connectivity in functional networks of excitatory and inhibitory neurons (**Fig. 6C, D**). The spike frequency of inhibitory neurons tended to be higher than that of excitatory neurons (**Fig. 6E**). Network models of organoid networks revealed that inhibitory neurons in all organoid conditions were correlated to numerous excitatory neurons (**Fig. 6D, S8F, G, and H**). In V247X organoids excitatory-inhibitory/inhibitory-excitatory connectivity exhibited reduced strength while excitatory-excitatory connectivity increased in relative strength compared to isogenic control organoids (**Fig. 6F**). In contrast, no significant differences for these types of connectivity were observed in R306C organoids (**Fig. 6F**). Ca^2+^ imaging was not able to specify the information flow direction, that is whether E-to-I or I-to-E connectivity was altered; however, scRNAseq and NeuronChatDB indicated that only I to E connectivity was altered, likely due to the downregulation of GABA_A_ receptors in excitatory neurons in V247X mutant organoids. Thus, different mutations lead to distinct influences on cell-specific connections.

**Figure 6.**
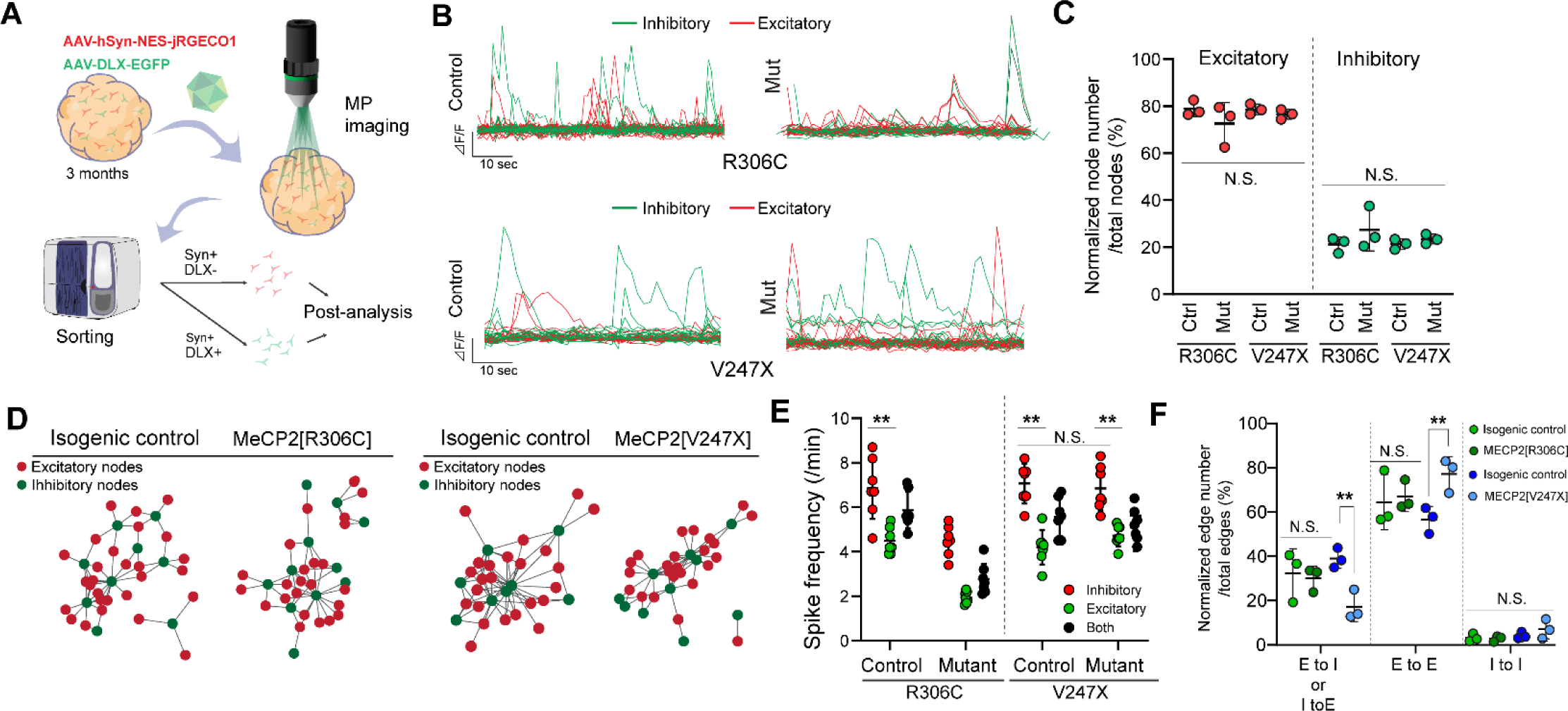
DLX labeling combined with Ca^2+^ imaging to determine neuronal activity in excitatory and inhibitory neurons and their networks. **(A)** DLX promoter-driven EGFP was co-expressed with jRGECO1a under hSyn promoter to label GABAergic inhibitory neurons 2 weeks prior to imaging. Time-lapse images were acquired by two-photon and three-photon microscopy. After imaging, the brain organoids were dissociated as single cells and sorted according to Syn+/Dlx-(Excitatory) and Syn+/Dlx+ (Inhibitory) neurons, followed by RNA extraction and PCR. (**B**) Ca^2+^ imaging of neuronal responses; neurons were clustered into excitatory and inhibitor neurons based on Dlx labeling in post-hoc analyses. (**C**) Quantification of normalized node number (the number of neurons) in all brain organoids. n = 3 organoids per condition. (**D**) Representative E/I networks. E/I tags were applied to graph theory analyses for plotting functional networks. Green dots indicate inhibitory nodes and red dots indicate excitatory nodes. (**E**) Separately calculated spike frequency in inhibitory and excitatory neurons based on labeling information. n = 6 organoids per condition. (**F**) Quantification of normalized number of edges for E/I nodes, E/E nodes, and I/I nodes defined from E/I networks. n = 3 organoids per condition. \**p*<0.05, \*\**p*<0.01; one-way ANOVA. Error bars indicate the ±SD.

## Discussion

In this paper, we describe altered functional connectivity of neurons and transcriptional changes in Rett syndrome patient-derived organoids carrying MeCP2[R306C] and [V247X] mutations. Our findings demonstrate that cerebral organoids can recapture developmental maturation of multiple neuron classes (**Figs. 1, 5**) along with their alteration by MeCP2 mutations. The two mutations that we examined, both in the TRD, led to some similar but importantly several different outcomes from morphological (**Fig. 1**), functional (**Figs. 2, 3**) and transcriptomic (**Fig. 5**) perspectives. V247X mutant organoids, but not R306C mutant organoids, were significantly larger and exhibited enhanced proliferation of neural progenitor cells (**Fig. 1**), confirming previous reports of V247X organoids showing expanded size and rosettes ^40^. We used a Ca^2+^ sensor together with three-photon microscopy to show that neurons in both R306C and V247X mutant organoids exhibited lower average spike frequency and reduced synchronicity compared to the isogenic pair, indicating disruptions in synaptic connections (**Fig. 2**). These data are consistent with previous reports of altered neuronal activity in RTT organoids ^26^. Graph theory-based functional network analyses demonstrated a greater number of connections per neuron in organoids with both RTT mutations (**Fig. 3**). These analyses revealed altered functional connectivity and network structures, characterized by changes in the local clustering coefficient and small world propensity (SWP); SWP was highly reduced in R306C organoids compared to controls (**Fig. 3**). Network analyses of human functional brain activity data has shown that neurons may be efficiently linked by local and long-range connections to represent robust and globally efficient processing^27^. Our findings demonstrate that even cerebral organoids have small world structure, indicating organized rather than random connectivity, and mutant organoids display disruption of small-world propensity. Additionally, third harmonic generation (THG) imaging allowed us to visualize the laminar structure of organoids alongside Ca^2+^ activity, demonstrating that intra-layer functional connectivity of neurons was more abundant than inter-layer connectivity in control organoids. However, this trend weakened in RTT mutant organoids, especially with the V247X mutation (**Fig. 4**).

Bulk RNAseq and scRNAseq results followed by NeuronChatDB analysis showed altered gene expression of HDAC2 in neurons in R306C organoids and impaired GABA-A receptor expression in excitatory neurons in V247X organoids (**Fig. 5**). Treatment with an HDAC2 selective inhibitor and a GABA agonist respectively in the two organoid mutations restored neuronal functionality, including spike frequency and SWPs (**Fig. 5N, O**). HDAC2 is an important transcriptional co-repressor with MECP2, and the R306C mutation has been shown to disrupt the interaction between another class 1 HDAC, HDAC3, and the NCoR interaction domain ^41^. This disruption may lead to abnormal HDAC2 gene expression^42^, which has previously been implicated in neurodegeneration and brain aging, such as Alzheimer’s disease ^43–45^. Restoration of HDAC2 levels has been shown to improve neuronal maturation and reduce amyloid-beta peptide along with restoring mitochondrial dynamics^46^. Although further assessment is necessary, our result suggest that selective restoration of HDAC2 could be one of the strategies for restoring phenotypes in R306C and other Rett syndrome mutations that target similar mechanisms. The V247X mutation causes upregulation of specific miRNAs, especially miR-199 and miR-214, in organoids^40^, by mechanisms that include aberrant BMP4 and SMAD signaling^9^. Local miRNA-dependent translational regulation of GABA-A receptor subunits has been described^47^, and could represent a mechanism by which V247X mutation-dependent miRNAs downregulate GABA-A receptors.

By labeling DLX-promoter driven inhibitory neurons with EGFP and using jRGECO1a as a Ca^2+^ indicator, along with fast-scan two-photon microscopy, we identified responses of inhibitory (DLX^+^) and putative excitatory (DLX^-^) neurons (**Fig. 6**). Excitatory-inhibitory connectivity exhibited reduced strength while excitatory-excitatory connectivity increased in V247X organoids compared to isogenic controls, whereas no significant differences in connectivity were observed in R306C organoids. Additionally, scRNAseq and NeuronChatDB analysis revealed reduction of I to E connection strength via downregulation of *GABRA2* and *GABRA4* in excitatory neurons and impairment of GABA-GABRA and NRX-NLGN signaling pathways, which may trigger altered network connectivity and small world propensity in V247X organoids. CellChatDB analysis revealed that astrocyte-to-excitatory neuron communication in V247X organoids was significantly deficient. Specific signaling pathways involving *NRG3-ERBB4, NRXN3-NLGN1,* and *Glu-SLC1A2/3-GLS to GRIAs* were disrupted, which are fundamental for astrocyte-neuron adhesion and glutamate regulation at the synapse^39,48^. These findings add to the evidence that astrocytes have critical and diverse roles in the pathophysiology of Rett Syndrome^49–51^, and our findings suggest that the effects may be mutation-specific.

Preliminary EEG data from human patients carrying truncating [R255X and R270X] or missense [R306C] mutations in the TRD indicated that SWPs were altered compared to age-matched controls in manner similar to changes observed in organoids with truncating (V247X) or missense (R306C) mutations in the TRD, suggesting that large-scale connectivity changes in Rett syndrome patients may originate in early development (**Fig. S4 and S5**). SWP derived from fMRI measurements is also disrupted in schizophrenia patients ^52^. Further investigation is essential to understand how changes in E-I balance associated with altered synaptic connectivity in MeCP2 mutations contribute to altered local and long-range networks in cerebral organoids and human subjects.

Our utilization of a cerebral organoid model, combined with multiphoton imaging using Ca^2+^ tracers and cell-specific labeling and transcriptomic analyses, demonstrates a valuable tool for studying the early development of neuronal activity and networks in neurodevelopmental disorders. These findings hold promise for understanding mutation-specific mechanisms of Rett syndrome, and developing targeted mutation-specific therapeutics

## Acknowledgments

We thank Taylor Johns and Alexandria Barlowe for technical assistance, Taylor Johns for editorial assistance, and members of the Sur lab for discussions and advice. This work was supported by NIH grants R01MH085802 and R01NS130361, MURI grant W911NF2110328, the Picower Institute Innovation Fund, and the Simons Foundation Autism Research Initiative through the Simons Center for the Social Brain to M.S.; and fellowships from the Japanese Society for the Promotion of Science and the Uehara Foundation to Y.O.

## Author Contributions

T.O. and M.S. conceived and designed the experiments. T.O. performed the experiments and T.O. and Y.O. analyzed the data. T.O. applied models to describe biological data and produced the analyses and figures. C.D. established organoid lines. D.K., A.L., M.F., and C.N. provided EEG data. T.O. and M.S. wrote the manuscript.

## STAR Methods

### Rett syndrome patient-derived iPS cells and isogenic controls

Rett syndrome patient-derived hiPS cells carrying the heterozygous MECP2[V247X] mutation (female, 25Y, 705G deletion, frameshift; very rare in RTT patients) and its isogenic control were obtained from the Coriell Institute (WIC07i-07982-4(MT) and WIC07i-07982-2(WT)). The isogenic control cell line had both wildtype and mutant alleles of MECP2, but the mutant allele that had the MECP2 mutation was X-inactivated in all of the mutant cells (Fig. S1A). Another Rett syndrome patient-derived hiPS cell line, which carries the heterozygous MECP2[R306C] mutation (female, 8Y, missense; 4–7% of RTT patients), and its isogenic control was obtained from the Coriell Institute (WIC05i-127-325(MT) and WIC04i-127-33(WT)). The isogenic control was established by site-directed genome editing by CRISPR/Cas9 to repair the mutant allele of MECP2, resulting in isogenic control cells having only wild-type alleles of MECP2 (**Fig. S1A**) ^53^. Healthy human iPS cells were also obtained from Coriell Institute (GM23279, female, 36Y) as a non-RTT related healthy control. All hiPS cells were maintained on Matrigel-coated 6-well plates in mTeSR Plus medium (STEMCELL Technologies) with 10 μM Y-23632 (Rock inhibiter, STEMCELL Technologies) for the first day after each passage and subcultured every 5–7 days using ReLeSR reagent (STEMCELL Technologies). They were characterized by immunostaining with pluripotent markers (OCT3/4, Nanog, TRA-1-60, and SSEA4), and MECP2 mutations were identified by Sanger sequencing at the DNA level (**Fig. 1B**), allele-specific PCR (**Fig. S1F and E**), western blotting (**Fig. S1F**) and immunostaining of protein expression levels (**Fig. S1G**). In addition, whole exome sequencing was performed (Germline Whole Exome Sequencing (WES) v6.0., 2 x 150 bp reads, Broad Institute) to determine the MECP2 mutation and confirm the absence of other mutations (**Fig. S1D and F**). SNVs and indels were characterized by mapping on the reference genome (GRCh37/hg19).

### RTT cerebral organoid formation and differentiation

To generate cerebral organoids, two mutant hiPS cell lines and isogenic control iPS cells on 6-well plates were dissociated into single cells with TrypLE express and then plated at 30,000 cells per well on U-bottom ultra-low attachment 96-well plates (Corning) with mTeSR Plus supplemented with 20 μM of Y-23632. After 24 h, the culture medium was replaced with neural induction medium (knockout DMEM-F12, 15% (v/v) knockout serum replacement, 1% (v/v) MEM-NEAA, 1% (v/v) GlutaMAX, 100 nM LDN-193189, 10 μM SB431542, and 10 μM XAV939) and changed every other day. After 10 days of culture, the culture medium was replaced with 1:1 mixture of knockout DMEM/F12 and neurobasal medium supplemented with 0.5% (v/v) N2 supplement, 1% (v/v) B27 plus supplement with vitamin A, 1% (v/v) GlutaMAX, 1% (v/v) MEM-NEAA, and 0.25 mg/ml (v/v) human insulin solution. This medium was changed every 2 days until 18 days of culture had elapsed. After 18 days, the organoids were transferred to flat-bottom ultra-low attachment 24-well plates (Corning) on a rocking shaker (120 rpm) and switched to a maintenance medium (Neurobasal medium supplemented with 1% (v/v) N2 supplement, 2% (v/v) B27 supplement with vitamin A, 1% (v/v) GlutaMAX, 1% (v/v) MEM-NEAA, 0.25 mg/ml (v/v) human insulin solution, 20 ng/ml BDNF, 200 mM ascorbic acid, 1 mM dibutyryl-cAMP (Sigma-Aldrich), and 1% (v/v) Penicillin/Streptomycin). After 30 days from differentiation, the culture medium was changed twice per week with BrainPhys medium supplemented with 2% (v/v) B27 supplement with vitamin A, 1% (v/v) GlutaMAX, 20 ng/ml BDNF, 20 ng/ml GDNF, and 1%(v/v) Penicillin/Streptomycin until the time of imaging. For the drug treatment experiments, organoids were treated with DL-2-Amino-5-phosphonopentanoic acid (AP-5, 100 μM), CNQX;100 μM, and bicuculine 50 μM just before Ca^2+^ imaging. For chronic treatment, organoids were treated with Baclofen 20 μM and HDAC2 selective inhibitor (CAY-10683, Santacruzamate A, 50 μM) for 72 hours prior to imaging.

### Virus infection

To visualize neuronal activity by fluorescent Ca^2+^ indicators (GCaMP6f driven by CAG promoter and jRGECO1a driven by hSyn) and to label GABA-ergic interneurons under mDLX enhancer element, the organoids were transfected using AAV1. pAAV.CAG.GCaMP6f.WPRE.SV40 was a gift from Douglas Kim & GENIE Project (Addgene #100836). pAAV.Syn.NES-jRGECO1a.WPRE.SV40 was a gift from Douglas Kim & GENIE Project (Addgene plasmid # 100854; http://n2t.net/addgene:100854; RRID:Addgene_100854). pAAV-mDlx-GFP-Fishell-1 was a gift from Gordon Fishell (Addgene plasmid # 83900; http://n2t.net/addgene:83900; RRID:Addgene_83900). Additionally, 2 μL of AAV solution was added to 500 μL of medium in 24-well plates containing one cerebral organoid at least 1-2 weeks before multi-photon imaging. Then, the maintenance medium was replaced with BrainPhys Imaging Optimized Medium supplemented with 2% (v/v) B27 supplement with vitamin A, 1% (v/v) GlutaMAX, 20 ng/ml BDNF, 20 ng/ml GDNF, and 1% (v/v) Penicillin/Streptomycin before imaging.

### Three-photon and two-photon microscope for calcium imaging

To capture Ca^2+^ dynamics of neurons deep within live organoids, three-photon excitation at 1300 nm was used for both GCaMP6f with label-free THG, or co-labeling of jRGECO1a/DLX-EGFP with label-free THG, with a Spirit-NOPA laser system (30-40 fs, Spectra-Physics). A 1300 nm wavelength laser was pumped and compressed from a 1045 nm Spirit One laser (300 fs, 400 kHz, 16W, Spirit, Spectra-Physics). The laser beams were scanned by a pair of galvo-galvo scanners (6215 H, Cambridge Technologies) to image the laser spot on the back aperture of the objective (Olympus 25x, 1.05 N.A.) using a pair of custom-designed scan and tube lenses. A pair of collection lenses for four photon-multiplying tubes (PMTs) collected the emitted signal from the cerebral organoids. The frame rate at 256×256 pixel resolution was increased up to 5 Hz for functional imaging or 0.9 Hz for structural imaging with 512×512 pixels. All of the fluorescent emission light was initially separated by a primary dichroic short-pass filter mirror (890 nm). GCaMP6f (EGFP) signals were detected using GaAsP photomultiplier tubes (H7422A-40, Hamamatsu, Japan) with a band-pass filter (Semrock, 540/50 nm). jRGECO1a was detected using GaAsP photomultiplier tubes (H7422A-40, Hamamatsu, Japan) and a bandpass filter (Semrock, 610/50 nm). THG signal was detected using a bialkali (BA) photomultiplier tube (R7600U-200) and a narrow-bandwidth bandpass filter (Semrock, 430/20 nm). The laser power was attenuated by a polarizer down to 10 mW for surface observation and 40-50 mW for deep-tissue observation. Automated image acquisition and control of the scanners and sample stage was carried out using ScanImage (Vidrio) ^54^. The organoids were placed on an x-y motorized stage and the objective lens was placed on a single-axis motorized stage (MMBP, Scientifica) to move it in the z-direction.

For some experiments, we used DLX-labeled neurons and Ca^2+^ imaging simultaneously, employing fast scanning. For these, we used a two-photon microscope employing a galvo-resonant scanner (Bruker) with Insight X3 laser (Spectra-Physics) at 1040 nm for jRGECO1a excitation and a MaiTai laser (Spectra-Physics) at 860 nm for EGFP excitation. Both lasers were aligned with the same light path and could be scanned up to 60 Hz sampling frequency at 512×512 resolution. The emission light was initially separated by a primary dichroic short pass filter mirror (890 nm) and the green and red signals were separated by a dichroic mirror (Semrock, 540/50 nm) and further filtered by bandpass filters (525/25 nm and 610/45 nm, respectively). Both signals were detected using GaAsP photomultiplier tubes (H7422A-40, Hamamatsu, Japan). The microscope and laser were controlled by Prarie View software (Bruker).

### Data processing of Ca^2+^ imaging data set

Time-lapse calcium imaging data was processed using Suite2p (suite2p, python==3.9) to identify active neurons. Regions of interest (ROIs) were defined to encompass individual neurons within the field of view. Baseline fluorescence (F_avg) for each ROI was calculated as the average of the bottom 10% percentile of fluorescence intensity values. The relative change in fluorescence over baseline (ΔF/F) was computed as (F - F_avg) / F_avg. A photobleaching correction was implemented by normalizing the ΔF/F trace using the slope obtained from the initial (F_start) and final (F_end) fluorescence values. In experiments involving DLX co-labeling and mosaic organoids, Suite2p was additionally employed to identify and export binary masks corresponding to DLX-positive neurons (identified by EGFP expression) and EGFP-mutant neurons (using a threshold of 0.25). This information, along with ΔF/F and spatial coordinates (x, y) of each neuron, was exported for further analysis. Exported data (delta F, x-y neuron positions, and DLX index) were loaded in MATLAB and further analyzed according to data type. The delta F/F value was deconvolved to identify single neuron activity, and then neuronal spikes were detected by using the “findpeaks” function in MATLAB. The spike frequency was calculated in a specific period of time (minutes). To calculate the undirected graph, network nodes were defined by deconvolved deltaF/F of Ca^2+^ signals and a continuous measure of association between nodes was estimated based on the correlation or connection probability between neurons. Correlation matrices were generated by compiling all pairwise associations between nodes and applying a threshold to each element of this matrix to produce a binary adjacency matrix^27^. Finally, the network parameters of interest in these graphical models of the organoid network were calculated and plotted as a structural network that retains geometric information and a functional network that retains topological information determined by the pairwise correlation between neurons. In functional networks, the locations of nodes were calculated by force-directed layout based on the Kamada-Kawai algorithm^29^ and Fruchterman-Reingold algorithm^30^. Functional networks were mainly used for the analysis of organoids (**Fig. 3 and 4**) and structural networks were utilized for EEG analysis (**Fig. S4 and S5**). For functional analysis, the geometric information (x-y neuron positions) and the layer clustered index (superficial, intermediate, and SVZ/VZ layer) were included. Layer indexes were identified by THG images which were taken at the same location as the FOVs. Equations to calculate key parameters, including Pearson’s pairwise correlation, local clustering coefficient, path length, and small world propensity are given below.

#### Pairwise correlation coefficient

To determine the functional connectivity between two individual neurons *x* and *y*, Pearson’s pairwise correlation coefficients *r_xy_* were calculated after shifting to zero mean based on the following formula:

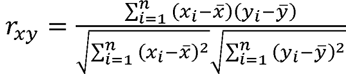

where *n* indicates number of time samples and *x* and *y* indicate the average response of neurons *x* and *y*.

#### Local clustering coefficient

The local clustering coefficient denotes the degree to which a vertex’s neighbors are connected to one another. We first calculated the local clustering coefficient c_i_ for each node i as:

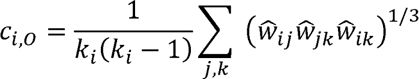

where w_ij_ corresponds to the strength of a connection between nodes *i* and *j*, = / ()^w w max w_ij_ w_ij_. w_ki_ correspond to the number of edges connected to nodes *i*. We thus defined the average clustering coefficient C, to be the average of the local coefficients:

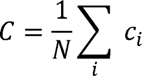

where, *N* corresponds to total number of nodes in the network.

#### Path length

Path length *L* is the average shortest distances between two nodes *i* and *j*, described as

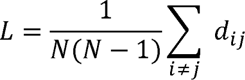

where, for a binary network, d_ij_ is the shortest path between nodes *i* and *j*. We defined the distance between two nodes as d_ij_ = 1/w_ij_.

#### Small-world propensity

To quantify the extent to which a network displays small-world structure, we defined the small-world propensity^55^, *ϕ*, to reflect the deviation of a network’s clustering coefficient, *C_obs_*, and characteristic path length, *L_obs_*, from both lattice (*C_latt_*, *L_latt_*) and random (*C_rand_*, *L_rand_*) networks constructed with the same number of nodes and the same degree of distribution: 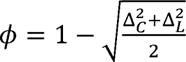,

where

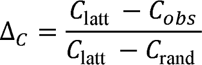

and

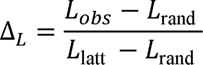

The ratios Δ_c_ and Δ_L_ represent the fractional deviation of the metric (*C_obs_*or *L_obs_*) from its respective null model (a lattice or random network). Networks with a high SWP can be categorized into three classes: (i) equally high clustering and low path length (both low Δ*_c_* and Δ*_L_*), (ii) high clustering and moderate path length (low Δ*_c_* and moderate Δ*_L_*) or (iii) moderate clustering and low path length (moderate Δ*_c_* and low Δ*_L_*).

### Determination of inter- and intra-layer connections

For label-free determination of the structure of brain organoids, a three-photon microscope and a THG channel were used, and the structural images were taken on top of functional imaging by GCaMP6f. Image analysis of the THG signal facilitated the extraction of individual neuron coordinates (x, y) within the 2D plane, thereby establishing their spatial distribution. Furthermore, the THG images were utilized to identify the center of rosette structures, a hallmark feature characteristic of the organoid surface. Subsequently, the angular relationship (θ) between each pair of neurons and the rosette center was calculated using the inverse cosine function (cos θ).

### Recording and processing of EEG data set

EEG data from Rett syndrome patients and typically developing subjects were recorded by the EGI NetStation system for data collection at a 1000 Hz sampling rate and downsampled to 256 Hz with 128 channels and analyzed a 10-20 electrode system for a recording period of 5 min. While recording, subjects sit in front of a monitor in a dimly lit room and watched videos screen savers-like video (as in the ABC-CT study ^56^). The recorded data were first processed with BEAPP to remove line noise and artifacts by ICA ^57^ and filtered with a 0.1 Hz high-pass filter and notch 60 Hz filter to remove DC noise. We analyzed 10 EEG resting recordings from 5 Rett syndrome patients and 5 typically-developed individuals, aged 3-22 years old. Two Rett syndrome patients had a MeCP2 truncating mutation (MeCP2[R255X]), two other patients had a truncating MeCP2[R270X]) mutation and another patient had an MeCP2 [R306C] SNP mutation. Processed time-series data were first segmented into 2 sec each for a total of 77 epochs, then power spectrum density was calculated for each of the 10-20 electrodes In addition, wavelet coherence between 1-100 Hz among all combinations of electrodes was calculated in each epoch and the calculated coherence was used as the weight of connections in the graph analysis. Based on wavelet coherence, topological analysis was performed and the degree of node, local clustering coefficient, and small-world propensity were calculated in the same way as for Ca^2+^ imaging data from organoids (**Fig. S4 and S5**).

### Microfluidic imaging chamber and temperature control system

To observe organoids in stable conditions under three-photon and two-photon microscopes, a microfluidic device was fabricated to fix the relative position of organoids in a reproducible manner during observation. SU-8 master positive patterned molds were fabricated by standard photolithography techniques described elsewhere ^58^. Briefly, SU-8 (2100) was poured onto a Silicon wafer (6-inches) and spin-coated (1200-1500 rpm, for 30 sec). The wafer was baked for 9 min at 65℃ and 40 min at 95℃ on a hotplate. Then, the wafer was irradiated by UV light (365 nm, 2.5-3.0 mWcm^2^,) with a photomask for 60-75 sec by Mask Aligner (EV620, TRL EV1). The wafer was baked for 7 min at 65℃ and for 13 min at 95℃ on a hotplate. After the wafer had cooled, SU-8 was developed using SU-8 Developer for 15 min, then washed with isopropyl alcohol (IPA) three times. The wafer was baked for 3 min at 150℃ in the oven. The thickness of the SU-8 was approximately 150 µm. A microfluidic device was made with a polydimethylsiloxane (PDMS) silicone elastomer kit Sylgard184 (Dow Corning). Silicone elastomer and a curing agent were mixed at a weight ratio of 10:1, degassed, poured onto the patterned SU-8 structures, and cured in the oven at 80 °C for 6 h. The holes for retaining cerebral organoids were created with biopsy punches (3 mm) and PDMS devices were sterilized by autoclave.

### Reverse-transcription PCR

To measure the biological activity of organoids, total RNA was isolated from tissues with TRIzol (Sigma) combined with Direct-zol RNA Purification Kits (Zymo Research). Reverse transcription was performed using a Superscript VILO cDNA Synthesis Kit (Thermo Fisher Scientific). The primer sequences are shown in **Table S2.** RT-PCR was performed with QuantStudio 3 (Applied Biosystems) using Takara TB Green Premix Ex Taq II (Takara). The mRNA level of glyceraldehyde 3-phosphate dehydrogenase (GAPDH) was used as the internal standard in all experiments. The RT-PCR experiments were repeated at least three times with cDNAs prepared from separate organoid pools.

### Flow cytometry and cell sorting

Single cells were dissociated by treatment with TrypLE Express (Thermo Fisher Scientific) or Gentle Cell Dissociation Reagent (Stem Cell Technologies) for 10-15 min. Then, the cell suspension was filtered with a 70-um cell strainer (BD Falcon) in 1% BSA-PBS. To sort inhibitory neurons and excitatory neurons, the cell population was gated by the hSyn-positive population and further gated and sorted as dlx-EGFP positive neurons (inhibitory neurons) and dlx-EGFP negative neurons (excitatory neurons) using BD FACSMelody (**Fig. S3**). After sorting, total RNA was collected for RT-PCR analysis. All other flow cytometry and analysis were performed with BD FACSMelody or FACSAria.

### scRNA-seq and processing of transcriptional data

Eight organoids in each condition were dissociated with TypeLE express for 5-10 min. The cells were then washed with PBS +0.1% BSA. Multiplexing oligos were incubated for 15 min and washed with PBS +0.1% BSA three times. The cell suspension was then pooled into one tube and used to perform 10X Chromium followed by sequencing with NovaSeq. Raw sequencing reads were processed using the 10x Genomics CellRanger bioinformatics pipeline v7.2. Cell ranger count matrices of four conditions (isogenic control/ MeCP2[R306C] organoid and isogenic control/ MeCP2[V247X] organoid) were integrated as Seurat objects in Seurat 5 (5.0.1) and Seurat wrapper (0.3.4) in R (4.3.2). Low-quality cells and doublets were removed from the following downstream analysis according to the number of genes (less than 500 and more than 7000) and mitochondria percentage (less than 20%). Combined count matrices were normalized and scaled and integrated with CCA algorithm and centered gene-wise, and then underwent dimensionality reduction using principal component analysis (PCA). After visual inspection of the PCA elbow plot, the top 12 PCs were chosen for further analysis. Clustering was performed on the PCs using the shared nearest neighbor algorithm in Seurat to obtain a UMAP plot. Clusters were identified by Louvain algorithm with multilevel refinement performed in Seurat. Unique cluster genes were identified using FindAllMarkers using the Wilcoxon rank-sum test without thresholds. To annotate the cell type, the following genes were used. RGCs cells: *VIM, HOPX, ASCL1, OLIG2, EOMES, ASPM*. Excitatory neurons: *DCX, BCL11A, BCL11B, GRIA1, GRIA2, GRIN2B, GRIN2A, FOXG1, FOXP2, SATB1, SATB2*. Inhibitory lineage: *DLX1, DLX2, DLX5, DLX6, GAD1*, *GAD2*. Astrocytes: *S100B*, *GFAP*, and *APOE* (**Fig. S4**). Based on these annotations, 5 different cell type populations were further clustered for sub-cluster analysis by recomputing the neighbor and dimensional reduction as well as UMAP plot for Pseudo-bulk RNAseq and pseudo-trajectory analysis. Single sample GSEAs were performed with escape R package (1.12.0) with Hallmark gene sets from MSigDB (https://www.gsea-msigdb.org/gsea/msigdb/). Pseudo-bulk RNAseq was performed by FindMarkers with sub-cluster and statistical significance was calculated by edgeR package (4.0.14) with Wilcoxon rank-sum test and visualized as a volcano plot. FDR was adjusted using the Benjamini-Hochberg method. Pseudo-trajectory was performed with the Monocle 3 package (1. 3. 4). To further characterize these clusters, bulk hallmark pathway analysis was carried out using the ClusterProfiler R package (4.10.1) and enricher with Hallmark gene sets and gene ontology gene sets from MSigDB. Ligand-receptor interaction analysis at the single cell level was performed using the CellChat R package (2.1.2) and NeuronChat R package. MicroRNA target predictions were performed based on miRPathDB (https://mpd.bioinf.uni-sb.de/). All cell ranger count matrices are deposited into the Gene Expression Omnibus as project number (https://www.ncbi.nlm.nih.gov/geo/000000000).

### Bulk RNA-seq and processing of transcriptional data

Cells from differentiated organoids (8 organoids) were lysed using TRIzol Isolation Reagent (Thermo Fisher Scientific) and kept at −80C until the next step. RNA was extracted and purified with Direct-zol RNA Purification Kits (Zymo Research). PolyA selection and stranded RNA-seq library preparations were performed with Illumina Stranded mRNA Pre-kit with rRNA depletion. RNA-sequencing was performed by the Broad Institute Sequencing Platform. 50M paired-end configuration and read-data were mapped to human reference genome GRCh37/hg19 or exosome sequenced genome of its cells. Mapped reads were quantified using featureCount. Differential expression was calculated by edgeR package in R, and used to determine significantly altered genes (adjusted p-value < 0.01).

### Statistical analysis

The reported values are the means of a minimum of three independent experiments. Data are presented as the mean ± SD. For equal variances and normality distribution, a Student’s t-test was performed. To compare groups at multiple conditions, statistical comparisons were performed using one-way analysis of variance (ANOVA), with post-hoc pairwise comparisons carried out using the Tukey-Kramer method. Statistical tests were performed using GraphPad Prism 9 (GraphPad Software, San Diego, CA). p-values <0.05 or p-values <0.01 were considered significant in all cases.

### Data availability

Read-level data from RNAseq and scRNA-seq will be deposited at GEO association. Any additional information required to reanalyse the data reported in this paper is available from the corresponding author upon request. The MATLAB for calcurating pairwise correlation of neuronal activity and graph theory to calcurate small workd proprensity and R code for RNAseq, scRNAseq is available from GitHub.

## Supplementary Figures and Tables

### This PDF file includes

Figures S1 to S8

Tables S1 to S2

**Fig. S1.**
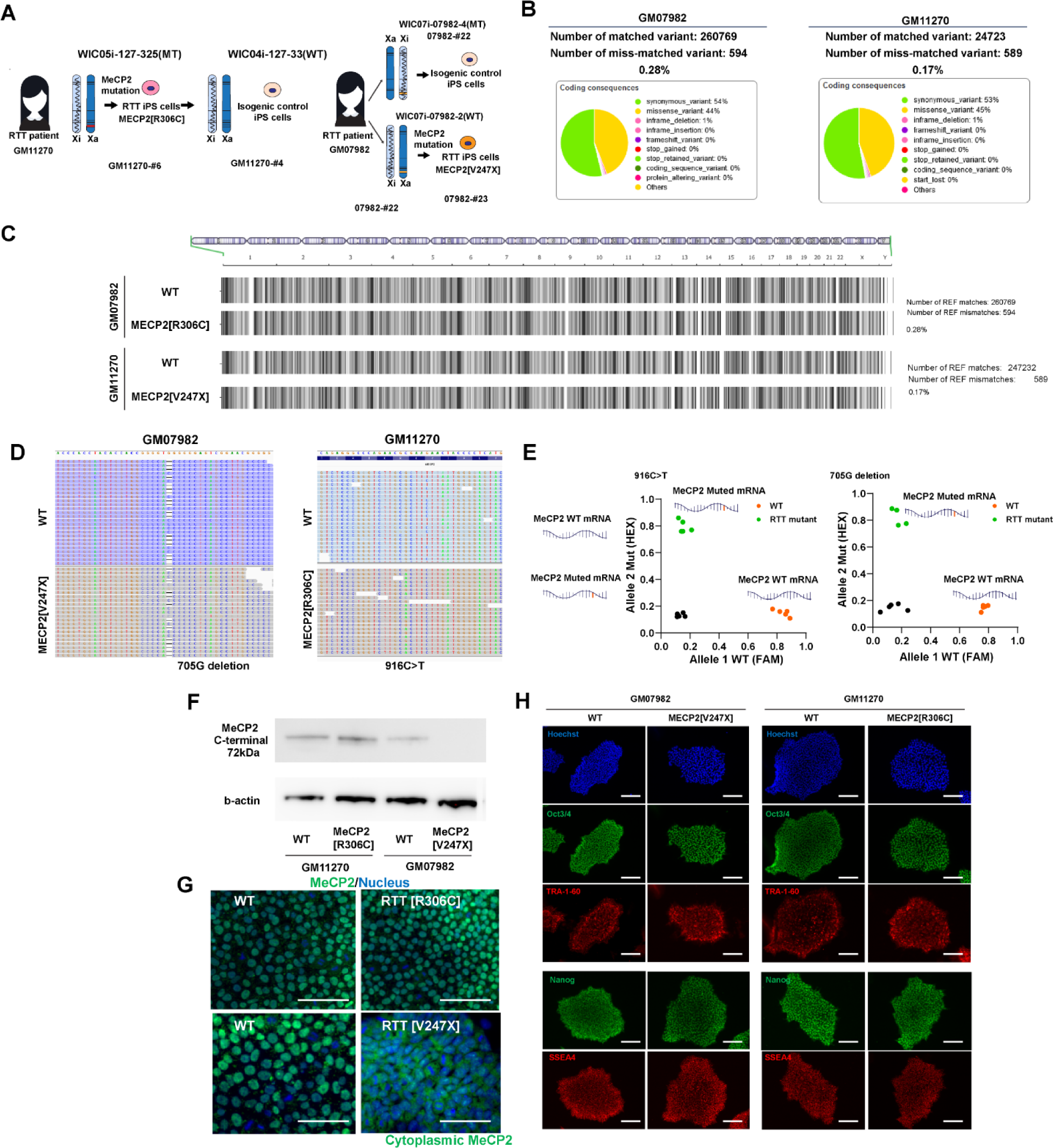
| Characterization of induced pluripotent stem cells (iPS cells) **(A)** Rett syndrome patient derived iPS cells and their isogenic controls, carrying MeCP2[R306C] and MeCP2[V247X] mutations. (**B, C**) SNV variations characterized by whole exosome sequencing revealed that 99.72% or 99.83% of genome was overlapped in isogenic control and mutant iPS cells; 0.28 and 0.17 mismatch was found in MeCP2[R306C] and MeCP2[V247X] lines respectively. (**D**) 705G deletion in MeCP2[V247X] mutant line and 916C>T conversion in MeCP2[R306C] mutant line. (**E**) Allele specific PCR in WT mRNA and mutant mRNA. (**F**) MeCP2 protein expression by western blotting. (**G**) MeCP2 expression by immunostaining. (**H**) Immunostaining for characterization of pluripotency by Oct3/4. TRA-1-60, Nanog, and SSEA4 in 4 iPS cells lines.

**Fig.S2.**
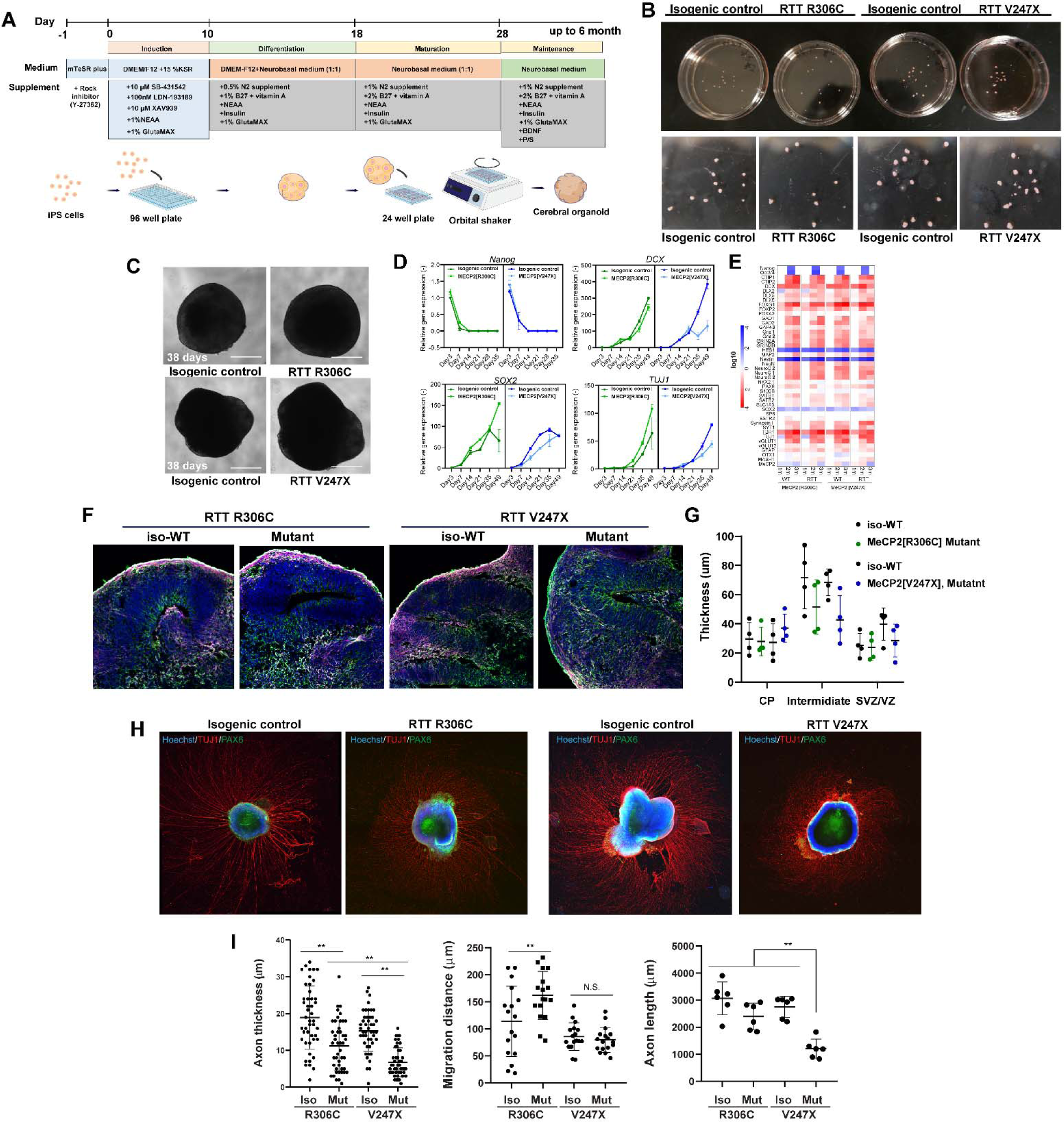
| Differentiation of cerebral organoids and their characterization. **(A)** Protocol for engineering cerebral organoids. (**B, C**) Representative images of cerebral organoids after 3 months of differentiation. (**D, E**) Characterization of brain organoids by RT-PCR during development (0-3 months). Expression of DCX and TUJ1, early neuronal markers, and SOX2, a neural stem cell marker, increased over time, whereas expression of Nanog, a pluripotency marker, decreased over time. n = 8 organoids. (**F**) Immunohistochemical staining to determine the layer structure of organoids. (**G**) Quantification of layer thickness (**I**) Immunostaining and axon elongation on Matrigel-coated plate. (**G**) Quantification of axon thickness, migration distance, and axon length. \**p*<0.05, \*\**p*<0.01; one-way ANOVA. Error bars indicate ±SD.

**Fig. S3.**
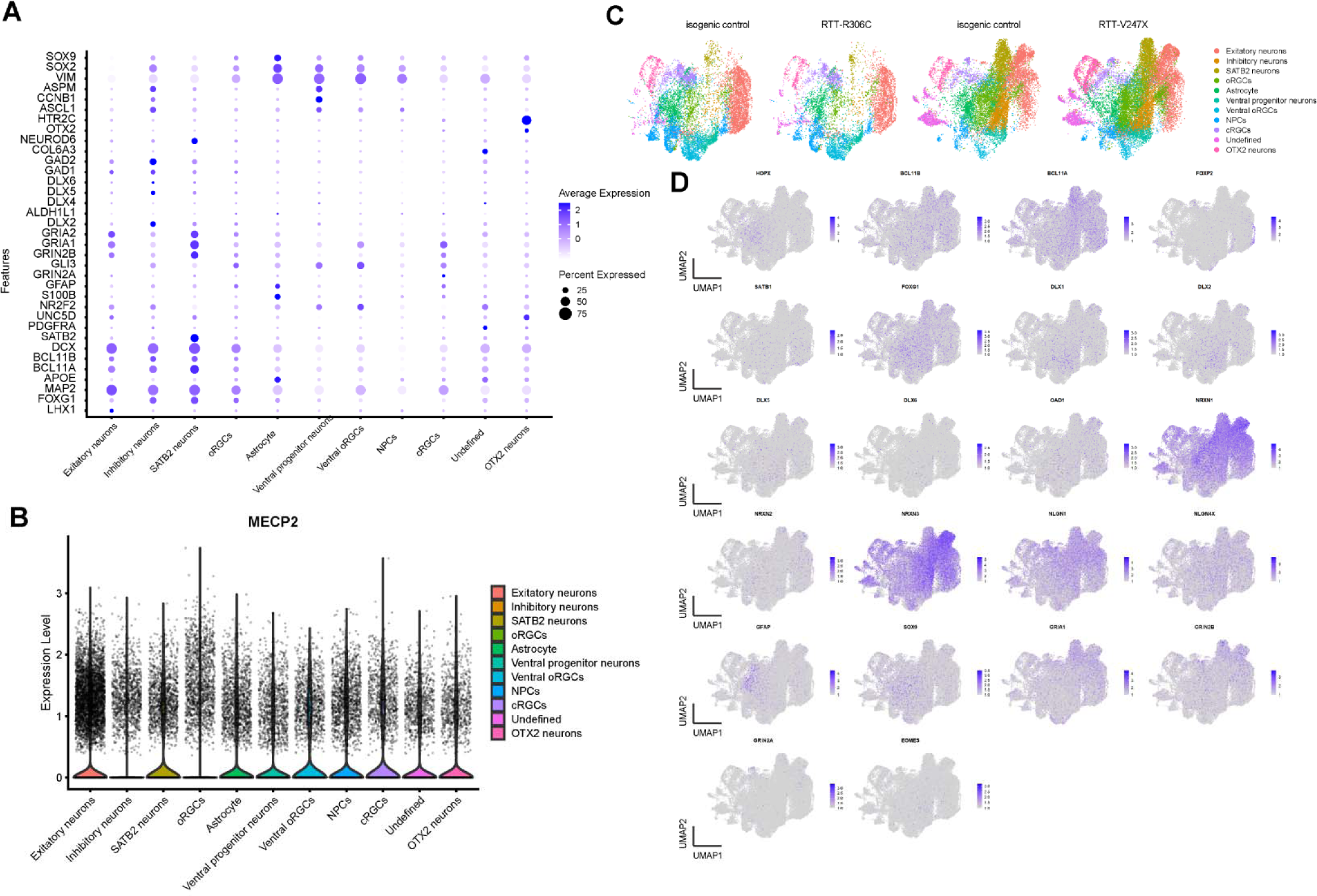
| Characterization of organoid development with scRNAseq. **(A)** Dot plot of cluster-defined feature genes used for annotation of cell types. (**B**) Violin plot of MeCP2 expression in different cell types clusters. No significant difference was seen among the cell-types. (**C**) Split view of dimplot of integrated data set. (**D**) Feature plot of the genes related to typical developmental processes used as an annotation of cell types.

**Fig. S4.**
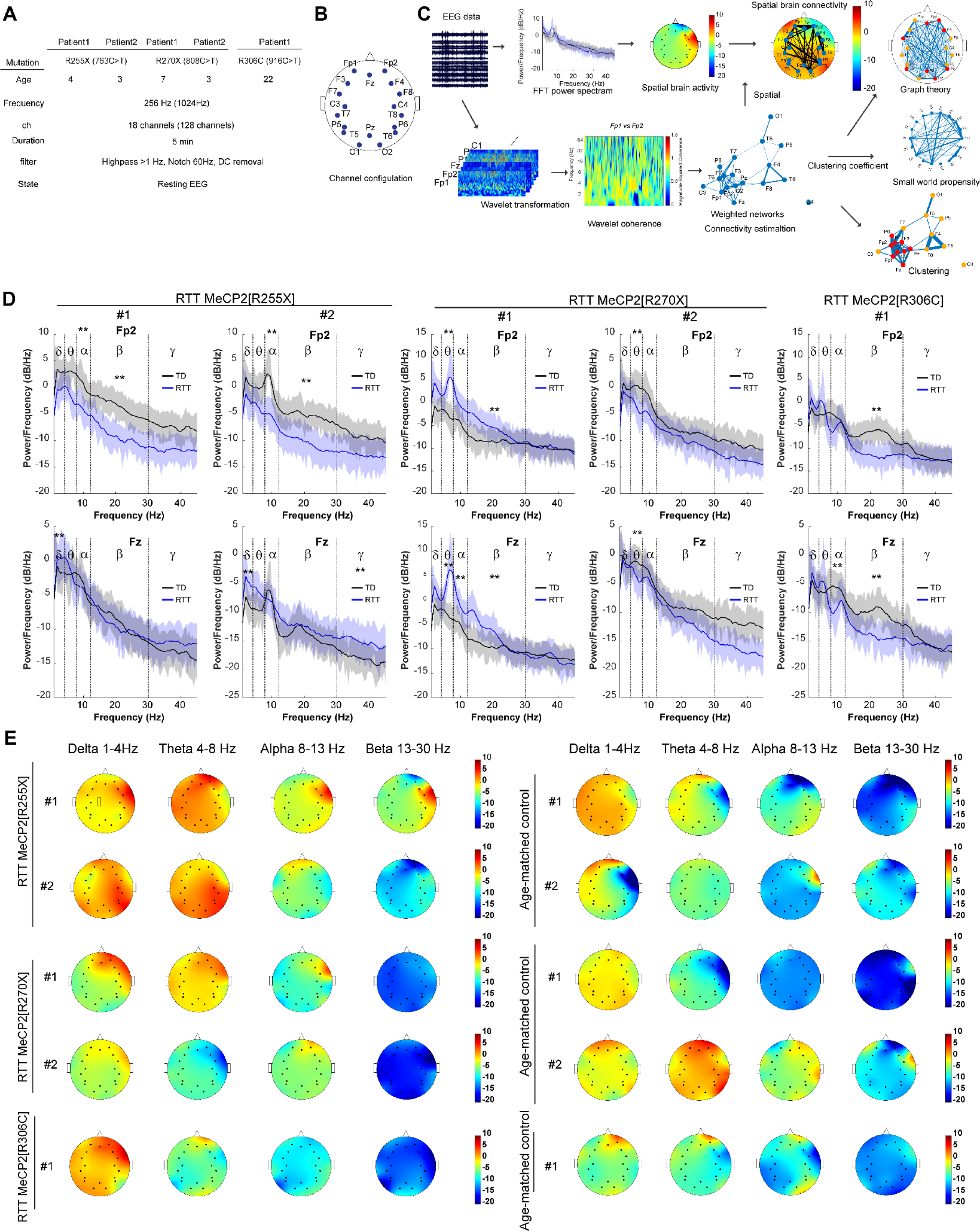
| EEG recordings from Rett syndrome patients and age-matched controls. **(A)** Patient information and EEG collection parameters for 5 Rett syndrome patients with MeCP2[R255X], MeCP2[R270X], or MeCP2[R306C] mutations and 5 age-matched controls. (**B**) EEG recording sites. (**C)** Analysis pipeline for EEG data. (**D**) Power spectrum density of EEG recording data between 0 and 45 Hz. (Blue: RTT patient: black: age-matched control). (**E**) Spatial brain activity in delta, theta, alpha and beta bands.

**Fig S5.**
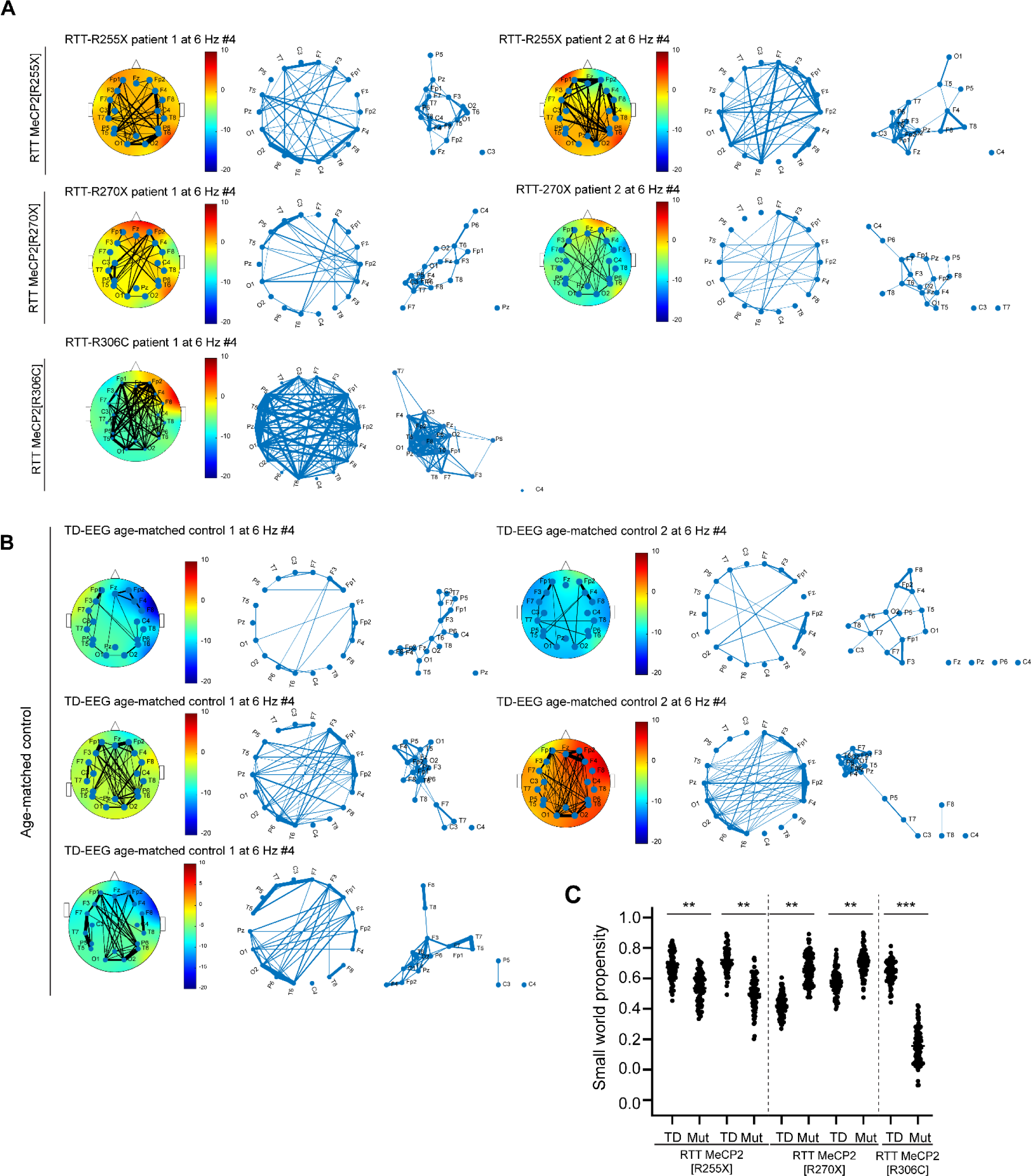
| Graph theory analyses of EEG data from Rett syndrome patients and age-matched controls. **(A)** Topological analysis of EEG data from 5 Rett syndrome patients, showing the pipeline of Fig. S4C. (**B**) Similar analysis of EEG data from age-matched controls. (C) Small world propensity comparing Rett patients and controls. \**p*<0.05, \*\**p*<0.01, *** p<0.001; one-way ANOVA.

**Fig. S6.**
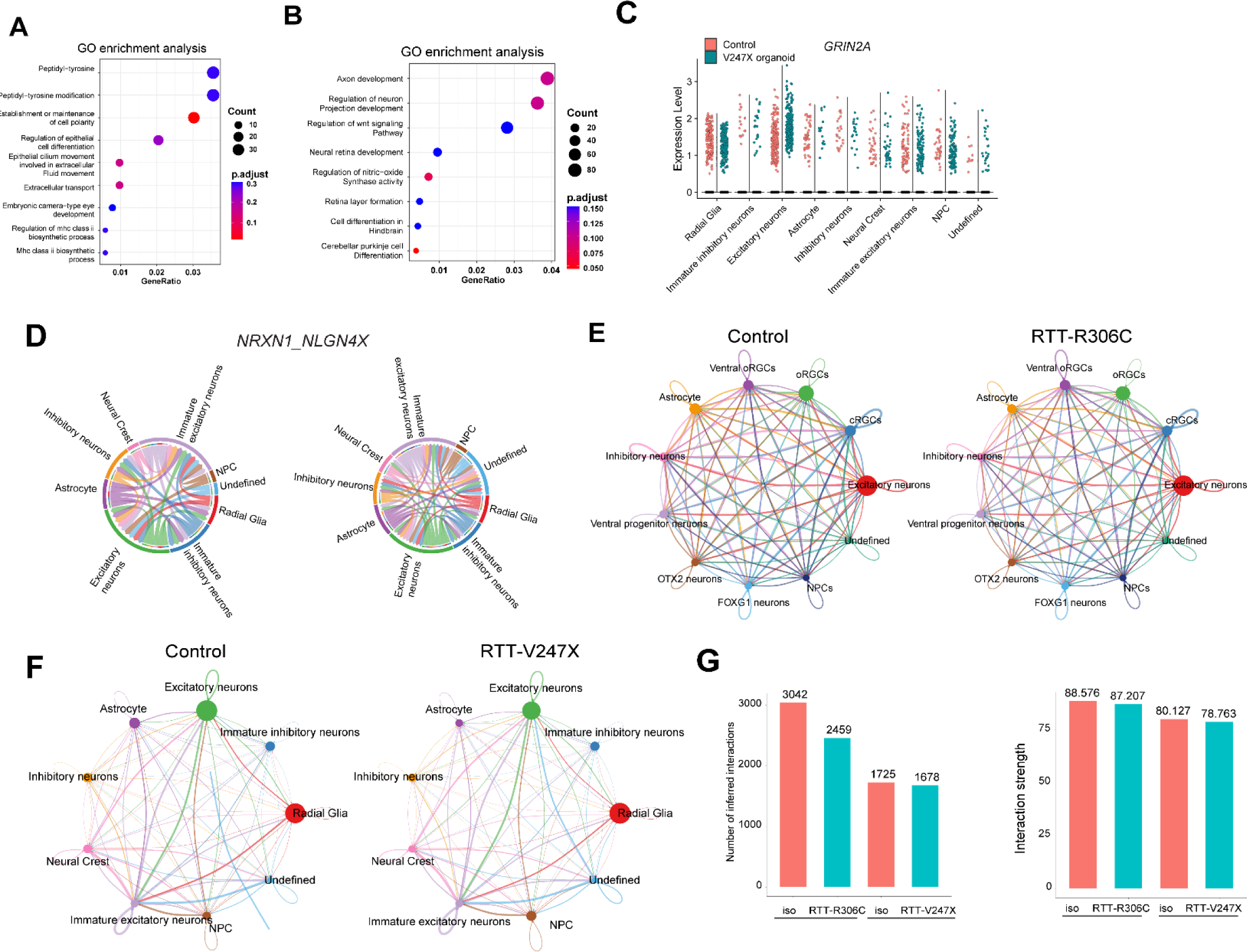
| Characterization of organoids with scRNAseq and CellChatDB. **(A)** GO enrichment analysis from downregulated genes in RTT-R306C mutant organoids relative to control (**B**) GO enrichment analysis from downregulated genes in RTT-V247X mutant organoids relative to control. (**C**) GRIN2A upregulation was observed only in excitatory neurons. (**D**) Chord plot showing the interaction between neurons and astrocytes via NRXN1_NLGN4X. (**E and F**) Analysis of cell-cell communication from scRNAseq data set with CellChatDB. Inferred inter-cellular signaling was visualized based on the number of links between the cells in the 4 conditions. (**G**) In RTT-R306C organoids, the total number of interactions in all the cell types was decreased compared to control, but not in RTT-V247X mutant organoids. The strength of communication was slightly decreased in both cases.

**Fig. S7.**
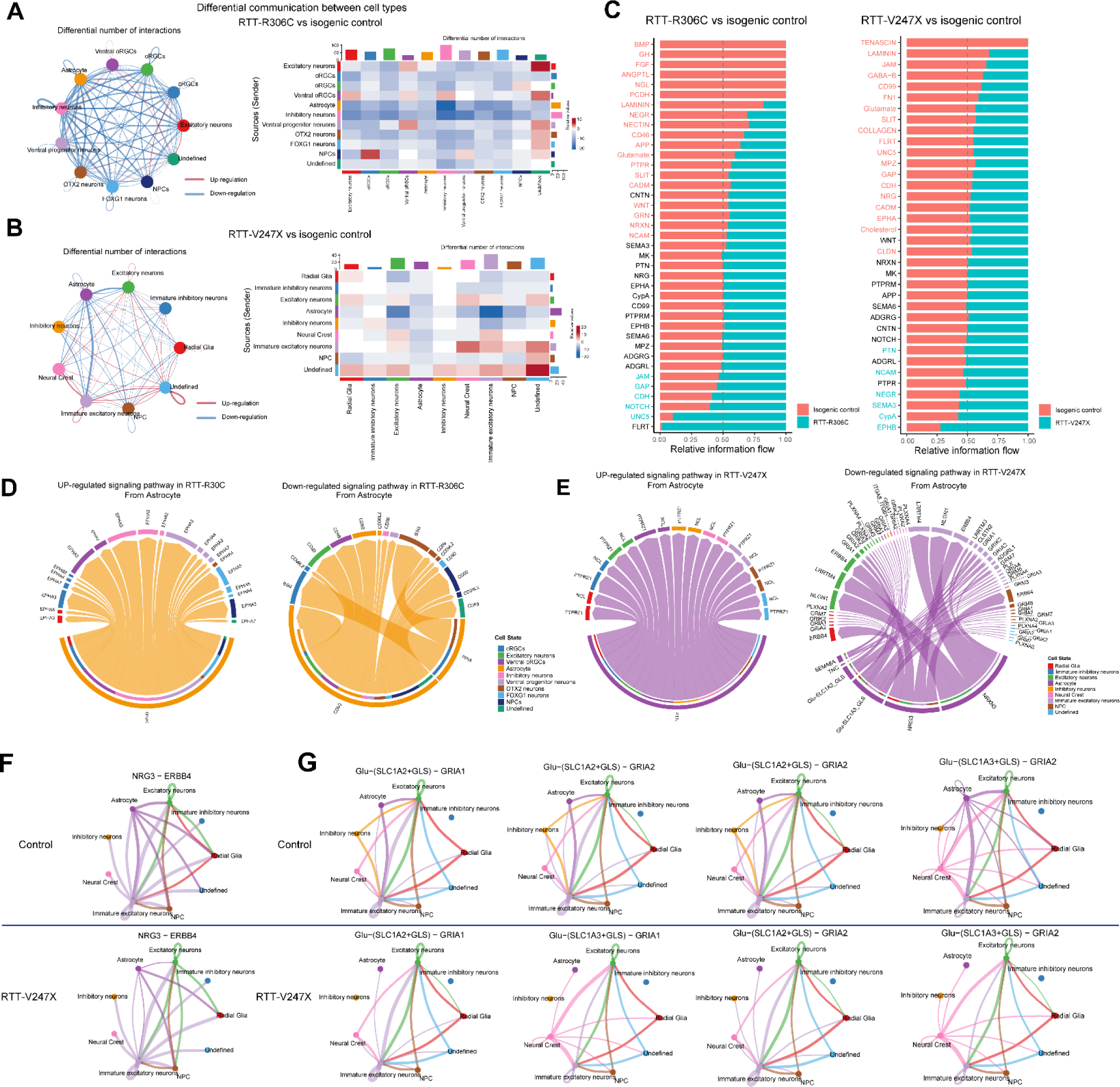
| Deficits in astrocyte communication identified by CellChatDB. **(A)** Differential cell-cell communication in RTT-R306C organoids vs control and RTT-V247X organoids vs control. Red: upregulated communication. Blue: downregulated communication. In RTT-R306C organoids, most intercellular communication was downregulated, and a few (NPC to oRGCs, ventral progenitor cells to ventral oRGCs, and ventral oRGCs to inhibitory neurons) were upregulated. (**B**) In RTT-V247X organoids, astrocyte-to-excitatory neuron and astrocyte-to-immature excitatory neuron communication were selectively deficient. (**C**) In RTT-R306C organoids, signaling pathways related to BMP, GH, ANGPTL, NGL, PCDH, LAMININ, NEGR, and NECTUIN were significantly downregulated, whereas TENASCIN, LAMININ, JAM, GABA-B, CD99, SLIT, NRG were downregulated in RTT-V247X organoids. (**D**) Circle chord plot showing all differential (upregulated and downregulated) intercellular signaling from astrocytes. In RTT-R306C organoids, CD99 (from astrocyte as sender) to CD99 (other cell type as receiver) and PPIA-BSG were determined as deficits in intercellular communication from astrocytes. (**E**) In RTT-V247X organoids, NRG3 (astrocyte)-ERBB4 (excitatory, astrocyte, and radial glia, and NPC), NRXN3(astrocyte)-NLGN1(excitatory neurons, immature excitatory neurons), Glu-SLC1A2/3-GLS to GRIAs intercellular communication were significantly altered. (**F, G**), Interaction plots for NRG and Glutamate pathways from astrocytes.

**Fig. S8.**
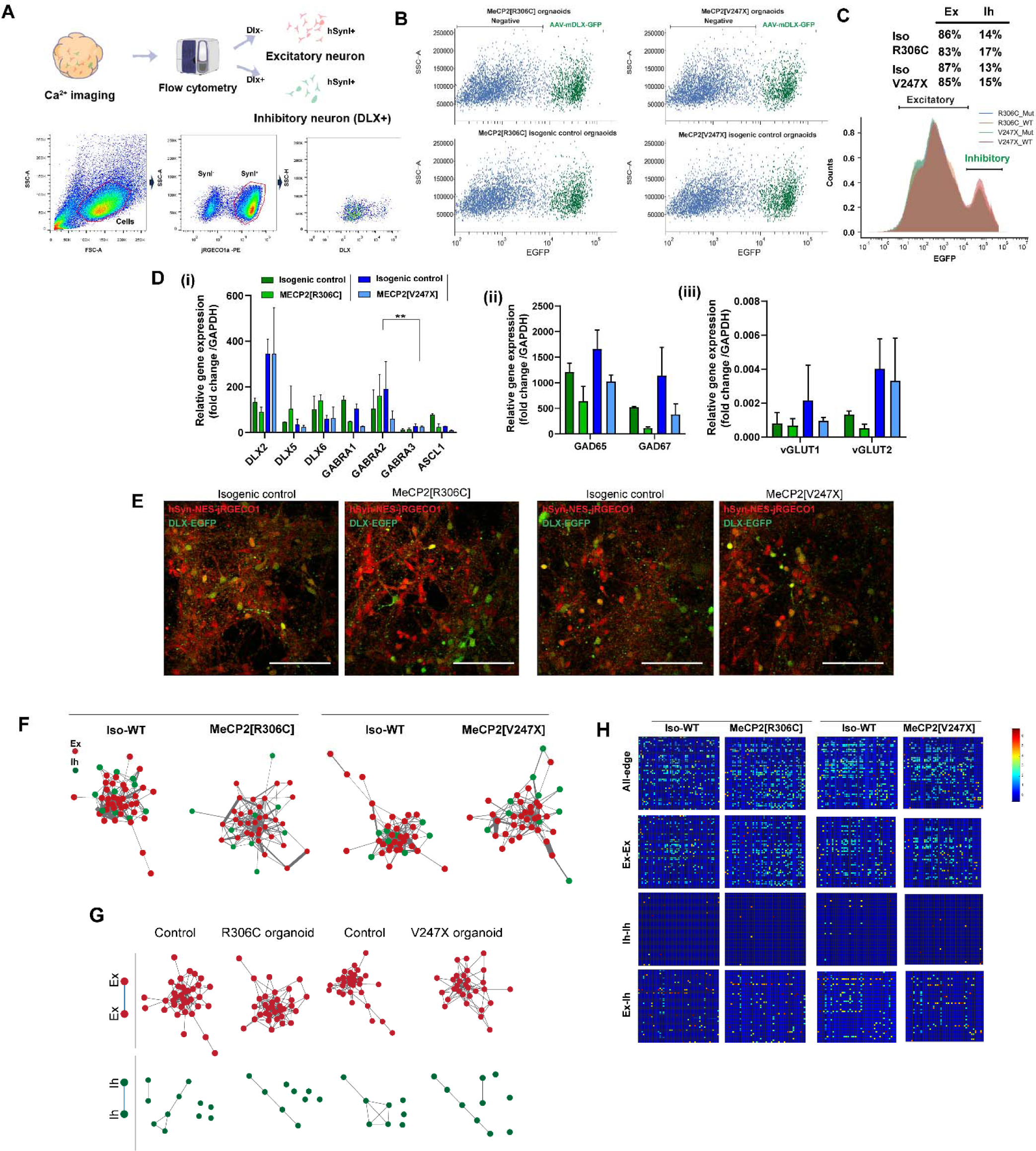
| DLX labeling of brain organoids and sorting-based analyses. **(A)** Sorting strategy of inhibitory and excitatory neurons. Inhibitory neurons were sorted as hSyn positive and Dlx positive cells. (**B, C**) Representative population of DLX+ and DLX-cells. (**D**) RT-PCR after sorting. (**E**) Representative images of neurons in 4 different brain organoids (R306C and V247X mutant and isogenic controls) labeled with DLX-EGFP and/or jRGECO1a. (**F**) Plots of functional networks in topological space with excitatory and inhibitory index labeling. (**G)** Edge connections between excitatory/excitatory nodes and inhibitory/inhibitory nodes extracted as subgraphs from all 4 types of organoids. (**H**) Correlation matrix, specifying Ex-Ex, In-In, and Ex-In nodes.

**Table S1.** Key resources.

| **Reagent and resource** | **Source** | **Identifier** |
| --- | --- | --- |
| **Cell line** |  |  |
| Rett syndrome patient derived iPS cells<br>(female, 25Y, 705G deletion) | Coriell Institute | WIC07i-07982-4(MT) |
| Isogenic control pair, wild type | Coriell Institute | WIC07i-07982-2(WT) |
| Rett syndrome patient derived iPS cells<br>(female, 8Y, missence) | Coriell Institute | WIC05i-127-325(MT) |
| Isogenic control pair, wild type | Coriell Institute | WIC04i-127-33(WT) |
| AAVpro 293T | Takara | N.A. |
| **Plasmid** |  |  |
| pAAV.CAG.GCaMP6f.WPRE.SV40 | Addgene | #100836 |
| pAAV.Syn.NES-jRGECO1a.WPRE.SV40 | Addgene | #100854 |
| pAAV-mDlx-GFP-Fishell-1 | Addgene | #83900 |
| pAdDeltaF6 | Addgene | #112867 |
| pAAV2/9n | Addgene | #112865 |
| pAAV2/1 | Addgene | #112862 |
| rAAV2-retro helper | Addgene | #81070 |
| **AAV** |  |  |
| AAV1. pAAV.CAG.GCaMP6f.WPRE.SV40 | Addgene | #100836-AAV1 |
| pAAV.Syn.NES-jRGECO1a.WPRE.SV40 | Addgene | #100854-AAVrg |
| pAAV-mDlx-GFP-Fishell-1 | Addgene | #83900-AAVrg |

**Table S2.**
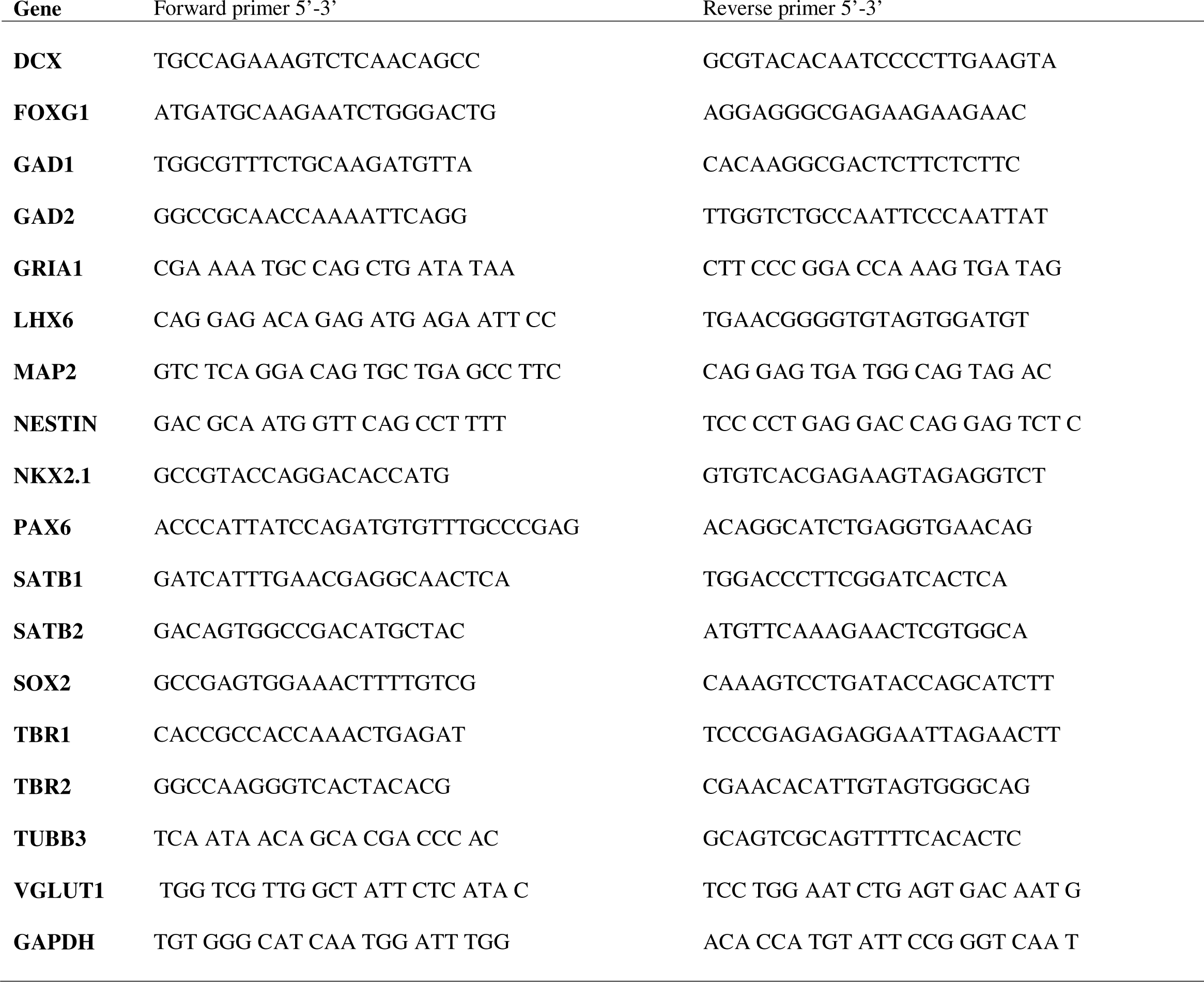
Real-time PCR primers.

